# Rqc1 and other yeast proteins containing highly positively charged sequences are not targets of the RQC complex

**DOI:** 10.1101/849851

**Authors:** Géssica C. Barros, Rodrigo D. Requião, Rodolfo L. Carneiro, Claudio A. Masuda, Mariana H. Moreira, Silvana Rossetto, Tatiana Domitrovic, Fernando L. Palhano

**Author notes:** These authors contributed equally (Géssica C. Barros, Rodrigo D. Requião, Rodolfo L. Carneiro).

## Abstract

Highly positively charged protein segments are known to result in poor translation efficiency. This effect is explained by ribosome stalling caused by electrostatic interactions between the nascent peptide and the negatively charged ribosome exit tunnel, leading to translation termination followed by protein degradation mediated by the RQC complex. These polybasic sequences are mainly studied in the context of artificial reporter systems. Examples of endogenous yeast proteins targeted by the RQC complex are Rqc1, a protein essential for RQC function, and Sdd1. Both contain polybasic sequences that are thought to activate the RQC, leading to protein down-regulation. Here, we investigated whether the RQC complex regulates other endogenous proteins with polybasic sequences. We show by bioinformatics, ribosome profiling data analysis, and western blot that endogenous proteins containing polybasic sequences similar to, or even more positively charged than those of Rqc1 and Sdd1, are not targeted by the RQC complex suggesting that endogenous polybasic sequences are not sufficient to induce this type of regulation. Finally, our results also suggest that Rqc1 is regulated post-translationally by the E3 component of the RQC complex Ltn1, in a manner independent of the RQC complex.

## INTRODUCTION

The first study to propose that the translation of positive amino acids could lead to translational stalling was published in 2008 by Lu and Deutsch [1]. It was already known by that time that the ribosomal exit tunnel, a narrow passage through which the nascent peptide passes as soon as it is synthesized, is negatively charged [2] and that the translation of the poly (A) tail from constructs lacking a stop codon, which leads to the incorporation of lysine residues (the codon AAA translates into lysine), causes translational repression [3-5]. This knowledge led to the hypothesis that a high concentration of positively charged residues inside the exit tunnel could slow translation rates due to the presence of electrostatic interactions. Using a cell-free approach, Lu and Deutsch, 2008, demonstrated that the presence of lysine or arginine repeats in a protein sequence is sufficient to decrease translation rates. They observed that the presence of as few as four nonconsecutive arginines in the construct was already enough to diminish the translational speed, while an equal concentration of negatively charged or uncharged residues did not affect translation rates. This phenomenon was also observed with lysines, and was independent of codon choice suggesting that the retardation effect was explicitly caused by the charge itself, not by the amino acid structure or tRNA availability [1].

These early experiments prompted the use of polybasic sequences as stalling regions in many subsequent papers. Most of these publications focused on the characterization of the ribosome quality control (RQC), a cellular pathway responsible for the co-translational degradation of nascent peptides [6, 7]. Polybasic sequences were added to or removed from a reporter protein, and the amount of translated protein was detected by western blot [6, 8, 9]. Using this approach, several authors have confirmed that the presence of polyarginine (R), polylysine (K), or mixtures of both amino acids can reduce protein production [10-12] and that this effect was linked to the action of the RQC machinery [6,13-16].

The RQC complex removes stalled nascent polypeptides from the ribosome and activates stress signals in *Saccharomyces cerevisiae* [6, 16]. A defective RQC complex causes proteotoxic stress in yeast cells [9] and neurodegeneration in mice [17]. The components of the yeast RQC complex are Rqc1, Rqc2, and Ltn1 proteins, all of which have a relatively low number of proteins per cell when compared with the number of ribosomes per cell, thus indicating that the system could easily reach saturation [6, 13, 18-20]. The RQC complex function is modulated by Asc1 and Hel2. These proteins detect ribosome stalling and promote early termination, leading to ribosomal subunit dissociation and transcript degradation [6]. After ribosome dissociation, the RQC complex can bind to the 60S subunit and induce the degradation of the nascent polypeptide through the E3 ubiquitin ligase activity of Ltn1 [6, 8]. Rqc1 and Ltn1 also recruit Cdc48 and its co-factors, Npl4 and Ufd1, which uses the energy from ATP hydrolysis to extract the stalled ubiquitylated polypeptide from the 60S subunit leaving the 60S subunit free to engage in another round of translation [7]. Rqc2 is involved in stress signaling through the formation of the C-terminal alanine-threonine (CAT) tail. After attaching to the 60S subunit, Rqc2 can incorporate alanine and threonine residues to the C-terminal of the stalled peptide, creating the CAT tail [9, 21-24]. This incorporation is random and independent of mRNA, and it is important to promote protein aggregation and/or to expose lysine residues that may be trapped inside the ribosome exit tunnel to the ubiquitin ligase activity of Ltn1 [23].

It has been proposed that the trigger for RQC activity is ribosome collision, with the formation of disomes, trisomes, and even tetrasomes [14, 25-28]. However, recent studies on the global analysis of disome formation suggested that collisions are relatively common, and most of these events are resolved without translation termination [29]. What determines whether a collision event will become a target of RQC or not is still an open question, but the disome structure, the time required to resume translation, and/or exposure of an empty A site, seem to play an important role in this decision [30]. Different studies have shown that the existence of stalling features in endogenous sequences seems to be accompanied by other unique characteristics, such as low initiation efficiency [31, 32] and/or the positioning of the stall sequence at the beginning of the transcript [33]. These adaptations are thought to result in fewer ribosomes per transcript, which would avoid ribosome collisions at the stalling-prone segments, thus preventing RQC activation.

In addition to the action as a salvage pathway for ribosomes stuck in unproductive translation events, it was suggested that the RQC pathway could regulate the endogenous expression of transcripts containing stalling features. This idea was put forward by an observation that Rqc1 levels increase with the deletion of *ASC1, HEL2, LTN1*, or *RQC2*, suggesting that Rqc1 protein expression could be controlled by RQC activity, providing negative feedback for the system [6]. Rqc1 contains a well-conserved K/R polybasic sequence in its N-terminal that, when substituted by alanine residues, led to increased protein expression. Based on these data, it was suggested that the RQC complex regulates Rqc1 protein levels through its polybasic sequence during Rqc1 translation [6]. More recently, the protein Sdd1, which has a polybasic sequence in its C-terminal, has been reported as another target of the RQC complex. Similar to Rqc1, the protein expression of Sdd1 was increased with the deletion of proteins of the RQC complex, or with mutations that substitute the positively charged residues from its polybasic sequence to alanine residues [25]. These observations led to the question of whether other endogenous proteins with similar polybasic sequences would be subject to the same regulation.

In this work, we first performed a bioinformatic screening to identify the yeast proteins containing polybasic sequences (sequences with the highest concentrations of K/R of the yeast proteome) Then, we analyzed previously published ribosome profiling experiments [25, 34-37] to access the impact of polybasic sequences on the translation of these proteins. Ribosome profiling is based on the deep sequencing of ribosome-protected mRNA fragments [38]. During translation, each ribosome encloses a portion of mRNA, protecting it against RNase digestion. This enclosure allows protected fragments to be sequenced, generating a map, at nucleotide resolution, of ribosomal occupancy on transcribed mRNAs. This map can report the occurrence of ribosome pausing or stalling, revealed by the accumulation of deep sequencing reads at specific positions of the transcript. We observed that polybasic sequences led to slower ribosome movement during translation and an increase in disome formation. However, we could not detect signatures of severe ribosome stalling, such as the presence of short (20-22 nt) mRNA fragments which indicates empty A site in ribosomes [39]. These data suggested that polybasic sequences of endogenous proteins do not lead to RQC complex activation. To test this hypothesis, we selected genes containing extremely positively charged segments and, without altering their original promoter or their 5’ UTR, we tested protein expression in yeast strains deleted for RQC components. In this experimental setting, we could not observe an increase in their steady-state protein levels, which is usually observed with reporters containing stalling sequences. These results suggested that RQC activity does not down-regulate translation of endogenous proteins with polybasic sequences. Our data indicate that polybasic sequences of endogenous proteins are sufficient to slow ribosome movement, but are not sufficient to induce the recruitment of the RQC complex. Rqc1 showed up-regulation in a *ltn1*Δ background but, in our hands, its levels were unperturbed when the complex was disabled at upstream steps (*asc1*Δ, *hel2*Δ) or at the CATylation step (*rqc2*Δ). Additionally, the deletion of *LTN1* increased the expression of the full-length Rqc1, suggesting that this regulation could not be coupled to a putative stalling event at a polybasic sequence located at the N-terminal. These results are inconsistent with the current model of Rqc1 being regulated by the RQC complex. Based on these data, we conclude that the RQC machinery is mostly dedicated to recycling ribosomes involved in aberrant mRNA translation and plays a small role in controlling endogenous protein levels under normal growth conditions. Moreover, we propose that Rqc1 regulation by Ltn1 is a post-translational event, independent of the RQC activity.

## MATERIALS AND METHODS

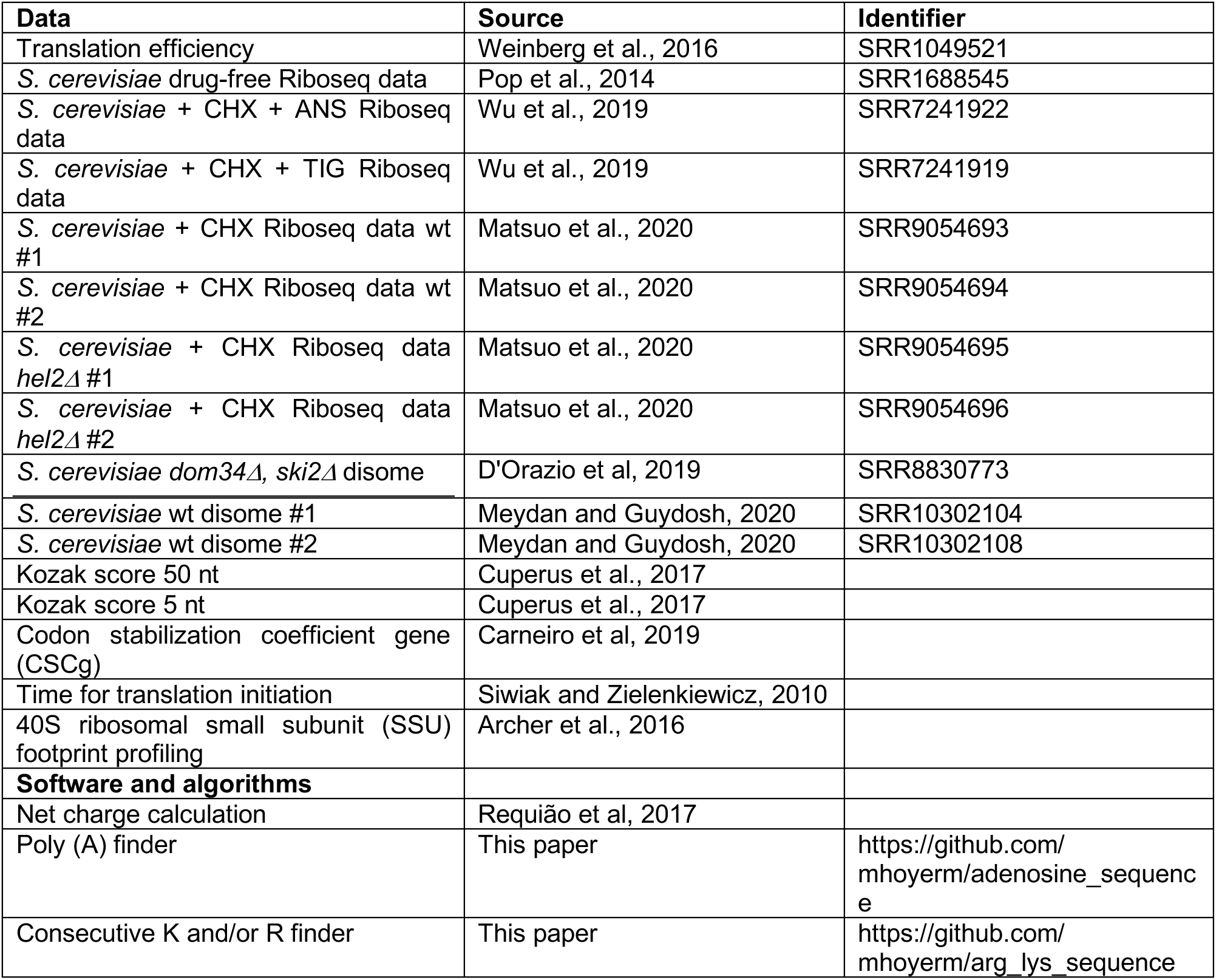
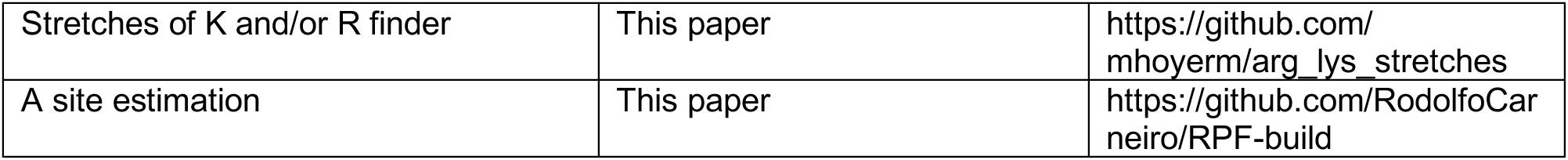

### Statistical analyses and raw data

The raw data used to create Figures 1 and 2, and the statistical analysis t-test calculations are presented in Supplementary Tables S1-2. The Spearman correlation among the parameters used in Figure 2 is presented in Supplementary Table S3. All statistical analyses were performed with GraphPad Prism 7.

**Figure 1.**
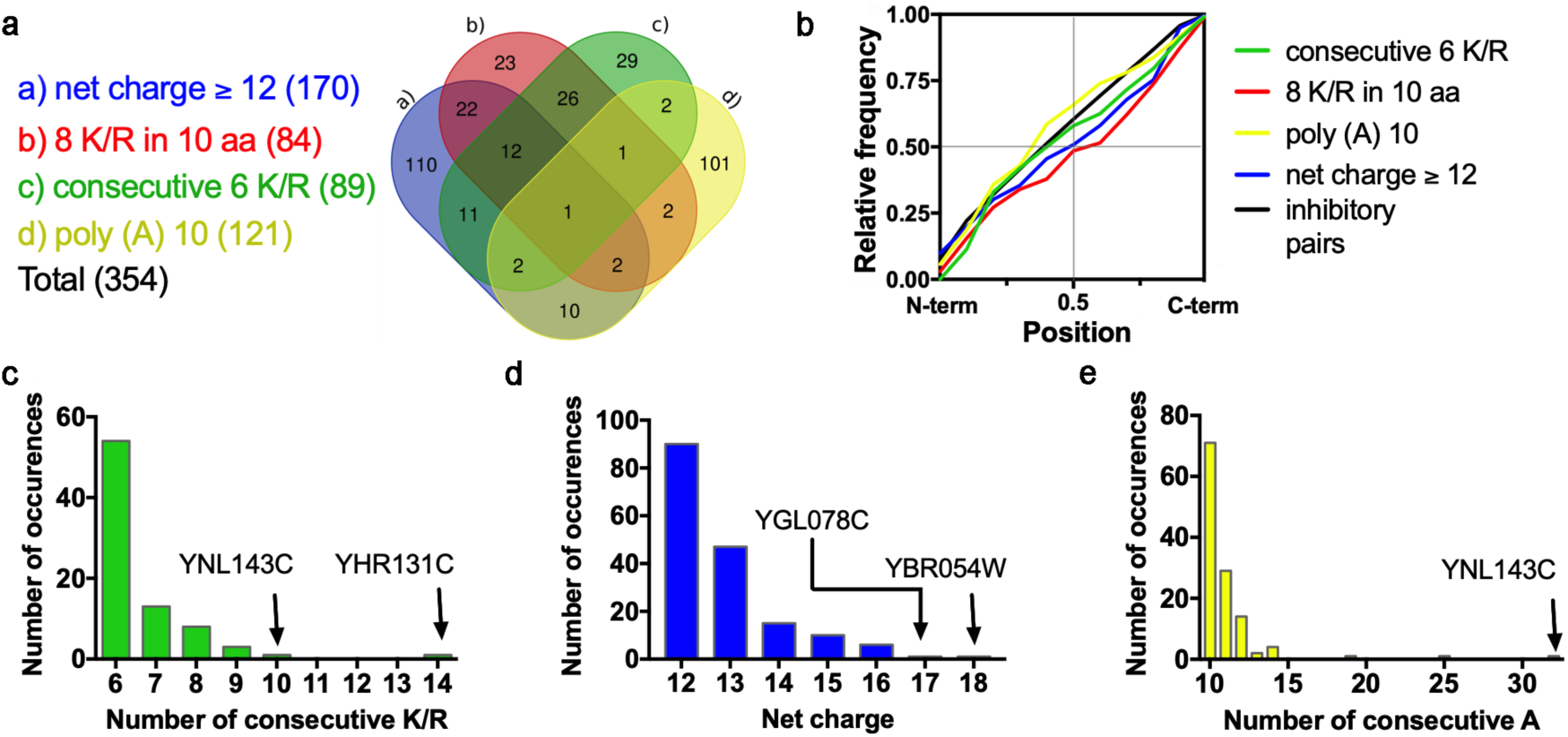
Bioinformatics screening to identify the polybasic sequences in the yeast proteome. (a) Venn diagram showing the genes identified as potential targets of the RQC complex. The list of identified genes is presented in Supplementary Table S1. (b) Cumulative frequency distribution of the location of the polybasic sequences of each group. For each gene, the desired attribute was located, and its position was determined in relation to the length of the gene. The frequency of distribution of the genes is depicted in the following categories: (c) consecutive 6 K/R, (d) net charge equal to or higher than + 12, and (e) 10 or more consecutive adenines. The genes with the highest scores of each category were highlighted.

**Figure 2.**
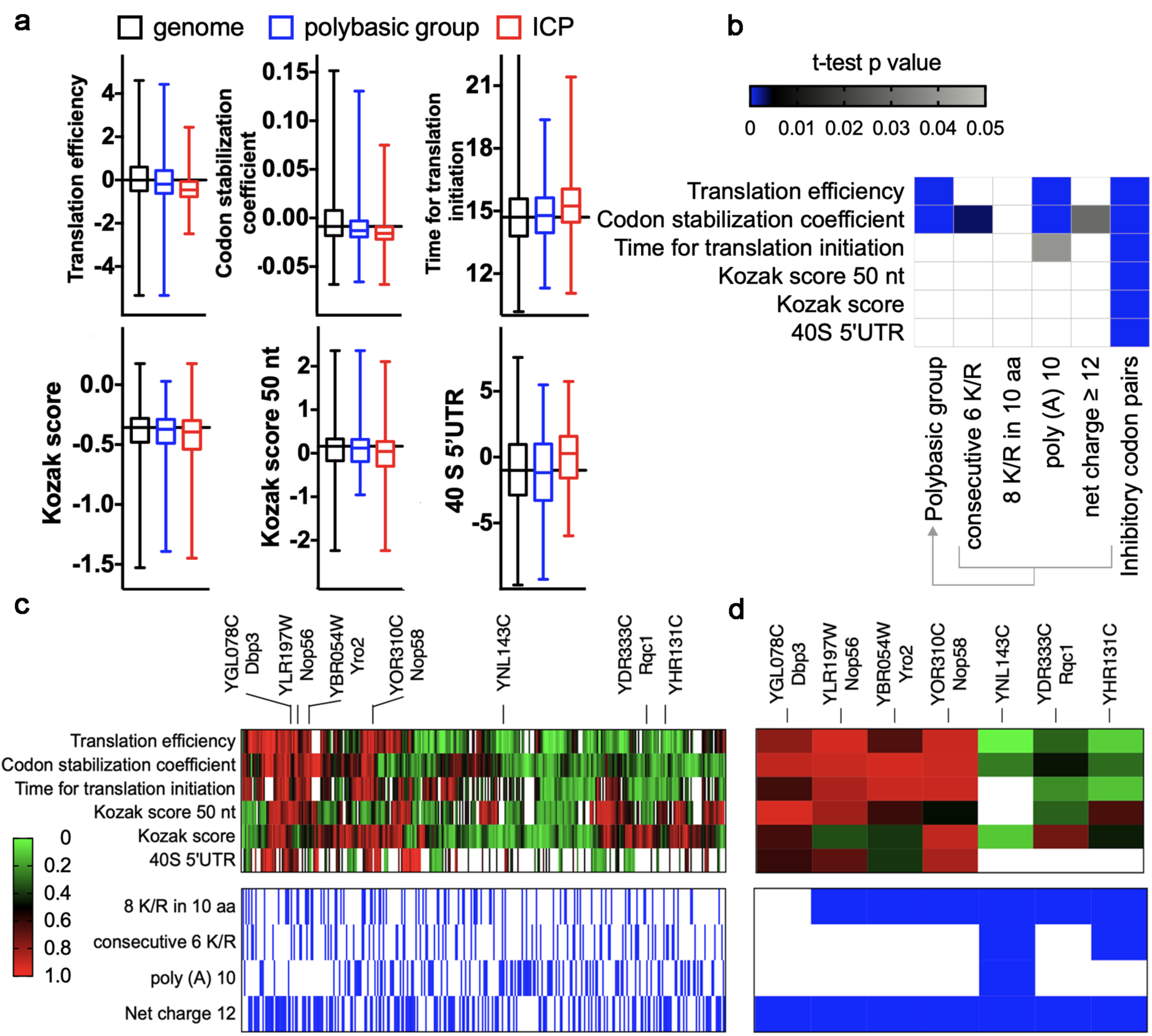
Translatability scores of genes with polybasic sequences. The different groups of genes with polybasic sequences were compared regarding parameters that, to some extent, reflect the efficiency of translation. (a) Box plot comparing different parameters from three groups of genes, namely, genome, polybasic group and, inhibitory codon pairs (ICP). (b) The Kolmogorov-Smirnov test p values are plotted for each comparison. The white rectangle means that a nonsignificant difference was found (p values >0.05). As a control, genes with one of the 17 inhibitory codon pairs (ICP) were used. (c) The values of the translation initiation parameters were normalized (where 0 and 1 represent the lowest and highest initiation rates of the full genome, respectively), and then the 354 identified genes were clustered by the Euclidean distance. The lower panel indicates the polybasic architecture present in each gene (marked in blue). (d) Translation initiation parameters cluster of the genes with the most prominent polybasic sequences (Figure 1c-e) and YDR333C/Rqc1 (included for comparison). The raw data, sample size, and p values are presented in Supplementary Table S2.

### Data sources

Coding sequences and the annotation of *S. cerevisiae* were obtained from the *Saccharomyces* Genome Database (SGD; https://www.yeastgenome.org) and Ensembl Genomes (http://ensemblgenomes.org/). We excluded dubious ORFs as defined in the *Saccharomyces* Genome Database from our analysis. The list of inhibitory codon pairs (ICP) and genes with ICP is presented in Supplementary Table S4.

### Net charge calculation

We developed a program that screens the primary sequence of a given protein and calculates the net charge in consecutive frames of a 30-amino-acid window, the approximate number of amino acids that fill the ribosome exit tunnel [40, 41]. For the net charge determination, K and R were considered +1; D and E were considered −1; and every other residue was considered 0. The additional charges of the free amino and carboxyl groups of the first and last amino acids, respectively, were disregarded. In a previous publication, we have shown that these simplified parameters are equivalent to a calculation using partial charges of individual amino acids at pH 7.4, according to their p*K*a values and the Henderson-Hasselbach equation [41]. The algorithm initially establishes a stretch containing a predetermined number of amino acids (#1 to #30, for example). The stretch charge is calculated, and the charge value and the position of the first amino acid are temporarily saved to the memory. Then, our algorithm advances to the next stretch, from amino acid #2 to amino acid #31, and performs the same analysis until the end of the protein.

### Codon stabilization coefficient (CSC) calculation

Based on Pearson’s correlation between the frequency of occurrence of each codon in each mRNA and the mRNA half-lives, Coller and colleagues created the codon occurrence to mRNA stability coefficient (CSC) [42]. We designed and implemented an algorithm that calculated the mean value of the CSC for each gene (CSCg) [43].

### Poly (A) finder

An algorithm was developed to locate the position of stretches containing only adenine (A) in an ORF. The algorithm finds adenine stretches of a chosen size for each ORF and generates as output a file containing the gene name, the stretch size, and its position in the ORF. For our analysis, we considered stretches of size equal to or larger than 10 adenines.

### Stretches of K and/or R finder

We developed an algorithm that searches for stretches containing a certain number of arginines and/or lysines in a stretch of 10 amino acids in a protein. The chosen number of K and/or R in the stretches was equal to or greater than 8. The input file of the algorithm was a proteome fasta file, and the output file returned the protein name and the position of the stretch found for each protein.

### Consecutive K and/or R finder

An algorithm was developed to find the location of stretches containing consecutive arginines and/or lysines in a protein. The stretches analyzed had a size equal to or larger than 6. The input file of the algorithm was a proteome fasta file, and the output file returned the protein name, the stretch size, and the position of the stretch in the protein.

### Yeast strains and growth conditions

Since no commercial antibodies are available for the proteins used herein, we used a TAP-TAG collection of yeast [44]. In this collection, each ORF was tagged with a C-terminal TAG, and the same antibody could be used to detect the full collection. For each potential RQC complex target analyzed, a knockout strain for the *LTN1* or *ASC1* gene was created by homologous recombination. For the Rqc1 TAP-TAG strain, knockout strains for *RQC2* and *HEL2* genes were also created. After the creation of the knockout strains, the level of the TAP-tagged proteins was compared with that of the wild-type strain by western blot analyses. The TAP-TAG collection strains were cultivated in YPD medium (1% yeast extract, 2% peptone, 2% glucose). The *S. cerevisiae* BY4741 strain transformed with GFP-R12-RFP vector (R12: CGG CGA CGA CGG CGA CGC CGA CGA CGA CGG CGC CGC) [6] was cultivated in minimum synthetic medium, uracil dropout (0.67% Difco Yeast Nitrogen Base without amino acids, 2% glucose, and 0.003% methionine, leucine, and histidine).

### Yeast strain constructions

For knockout creation, *S. cerevisiae* BY4741 yeast (TAP-TAG collection) was transformed using the PEG/Lithium Acetate method [45]. The PCR products used in the transformation were obtained by the amplification of the KanMX6 gene contained in the pFA6a-KanMx6 vector. All genes were knocked out by the insertion of the geneticin resistance gene into their original ORFs. The knockouts were confirmed using four sets of PCR. Two PCR sets combined primers for the target gene flank region (5’UTR or 3’UTR) with resistance gene primers. The other two PCR sets combined primers for the target gene flank region and ORF gene target primers. All the primer pairs used in the construction and confirmation of the mutants are described in Support Information.

### Protein extraction and SDS-PAGE

The protein extraction of the strains was conducted in the exponential phase of growth (O.D ∼ 0,6-0,8), treating equal quantities of cells using the NaOH/TCA method [46]. For the SDS-PAGE, equal sample volumes were applied on the acrylamide gels and submitted to an electrophoretic run. For the YHR131C TAP-TAG strain, protein extraction was conducted using a glass bead protein extraction protocol [47]. Total protein quantification of the glass bead protein extracts was conducted using the BCA Protein Assay kit (Thermo Fisher), and samples with equal amount of proteins were applied on SDS-PAGE gels. The acrylamide gel concentration changed between 10 and 12.5% according to the size of the analyzed protein.

### Western blotting

Proteins separated by SDS-PAGE were transferred to PVDF membranes using the TRANS-BLOT semidry system from BIO-RAD in transfer buffer (25 mM Tris, 192 mM glycine, 0.1% SDS, 20% methanol, pH 8.3). The membranes were blocked using a Li-Cor blocking buffer with an overnight incubation at room temperature. Incubation with primary antibodies was performed at 4°C for at least 1 hour. The membranes were incubated in Li-Cor Odyssey secondary antibodies at room temperature for at least 1 hour and then revealed in an Odyssey scanner. The primary antibodies used were mouse anti-PGK1 (Invitrogen; dilution 1:10,000), rabbit anti-TAP-Tag (Thermo Scientific; dilution 1:10,000), mouse anti-GFP (Sigma; dilution 1:2,000), and rabbit anti-Asc1 (dilution 1:5,000) [48]. The secondary antibodies that were used were anti-mouse 680 LT Li-Cor (dilution 1:10,000) and anti-rabbit 800 CW Li-Cor (dilution 1:10,000).

### qRT-PCR

For qRT-PCR analysis, the Rqc1 TAP-TAG knockout strains constructed in this work were cultivated until the exponential phase (O.D ∼ 0,6-0,8) and RNA from these cells were extracted following the hot phenol method. The same protocol was used for the corresponding knockouts from a commercial collection as a control. After extraction, the RNAs were submitted to DNase I (Thermo Scientific) treatment for DNA contamination elimination. The reverse transcription was performed using random primers with the High-Capacity cDNA Reverse Transcription Kit (Thermo Scientific). For the cDNA amplification, we used SYBR Green PCR Master Mix on the StepOne Plus System (Applied Biosystems). The following protocol was used on the thermocycler: 95°C for 10 min and 40 cycles of 95°C for 15s and 60° C for 1 min, melting curve analysis: 95°C for 15s, 60°C for 1 min and 95°C for 15s. The primers used were: 5’CGGAATGCACCCGCAACATT and 5’CGGCCTTTCCCATAATTGCCA for *RQC1*, and 5’ TTCCCAGGTATTGCCGAAA and 5’ TTGTGGTGAACGATAGATGGA for *ACT1.* The analysis of differential expression was made by relative quantification using 2^−ΔΔ**Ct**^ method, and the statistical analysis was conducted on Prism.

### Cycloheximide (CHX) treatment

For the experiments using CHX, concentrations ranging from 10^−6^ to 10^−1^ mg/ml were used, with concentration steps of 10 times the dilution factor. Strains grew until the exponential phase and were treated with CHX for 10 hours under shaking at 28°C.

### Ribosome profiling data

*S. cerevisiae* ribosome profiling data were treated as described previously [41, 49]. The data were analyzed as described by Ingolia and collaborators [38], except that the program used here was Geneious R11 (Biomatter Ltd., New Zealand) instead of the CASAVA 1.8 pipeline. The data were downloaded from GEO [34, 35], and the adaptor sequence was trimmed. The trimmed FASTA sequences were aligned to *S. cerevisiae* ribosomal and noncoding RNA sequences to remove rRNA reads. The unaligned reads were aligned to the *S. cerevisiae* S288C genome deposited in the *Saccharomyces* Genome Database. First, we removed any reads that mapped to multiple locations. Then, the reads were aligned to the *S. cerevisiae* coding sequence database, allowing two mismatches per read. For 27-29 nt footprint analyses, the ribosome profiling data were obtained without any drug, and just fragments with 27-29 nt were used [34]. For 20-22 nt footprint analyses, the ribosome profiling data were obtained with cells lysed with buffer containing 0.1 mg/ml of cycloheximide (CHX) and tigecycline (TIG) or 0.1 mg/ml of cycloheximide (CHX) and anisomycin (ANS) [35]. We normalized the coverage within the same transcript by dividing each nucleotide read by the sum of the number of reads for each gene in a given window. A window of 150 nucleotides before and after the beginning of a stretch of interest was used to calculate the number of reads of each gene. For Figures 3c-g, 5c, and Supplementary Figure S2, the ribosome profiling analysis was done following as described by Ingolia and collaborators [38]. The datasets used in our analyses were from Pop *et al.*, 2014 [34] (GEO accession GSE115162) and from Matsuo *et al.*, 2020 [25] (GEO accession GSE131214). The data were mapped to the R64.2.1 S288C Coding Sequence (CDS) from the *Saccharomyces* Genome Database Project. The alignment was done using Bowtie accepting two mismatches at most. The analysis was restricted to 28–32 nt reads for Supplementary Figure S2. The A-site was estimated based on the read length: it was defined as the codon containing the 16th nt for 28 nt reads; the 17th nt for 29, 30 and 31 nt reads; and the 18th nt for 32 nt reads, considering the first nt as 1. The A-site estimation and other Ribosome Profiling Analysis were performed by custom software written in Python 3.8.2. For Supplementary Figures S2 and S3, we compared the ribosome profiles normalized for genes of the five distinct groups (Supplementary Figure S2) or all polybasic group (Supplementary Figure S3) with 2,000 aleatory genes. Statistical analyses were performed with multiple t-tests using the Holm-Sidak method, and the adjusted p-value was calculated for each point (GraphPad Prism 7). For Figure 4, the ribosome profiling of each gene was downloaded from Trips-Viz by aggregating all experiments in the database [50]. Disome Profiling Data: The datasets we used for disome profiling were from D’Orazio et al, 2019 [36] (GEO accession GSE129128), and Meydan and Guydosh, 2020 [37] (GEO accession GSE139036). For the Meydan and Guydosh data, we further trimmed 2 nucleotides from the 5’ end of each read (the ones with the lowest quality in every read), and also 5 nucleotides from the 3’ end (adapting for the post-mapping trimmings done to the non-mapped reads, as described in Meydan and Guydosh, 2020). Non-coding RNA sequences were removed from both datasets using Geneious. Then they were also mapped to the R64.2.1 S288C Coding Sequence (CDS), using Bowtie, accepting two mismatches at most. For D’Orazio data, we accepted any read containing between 50nt and 80nt. We estimated the A site from the first ribosome (the one at the 3’ portion of the read) as the codon containing the 46th nucleotide. For Meydan and Guydosh data, we accepted reads containing between 57nt and 63nt. Also, we estimated the A site from the ribosome at the 3’ portion of the read as the codon containing the 46th nucleotide.

**Figure 3.**
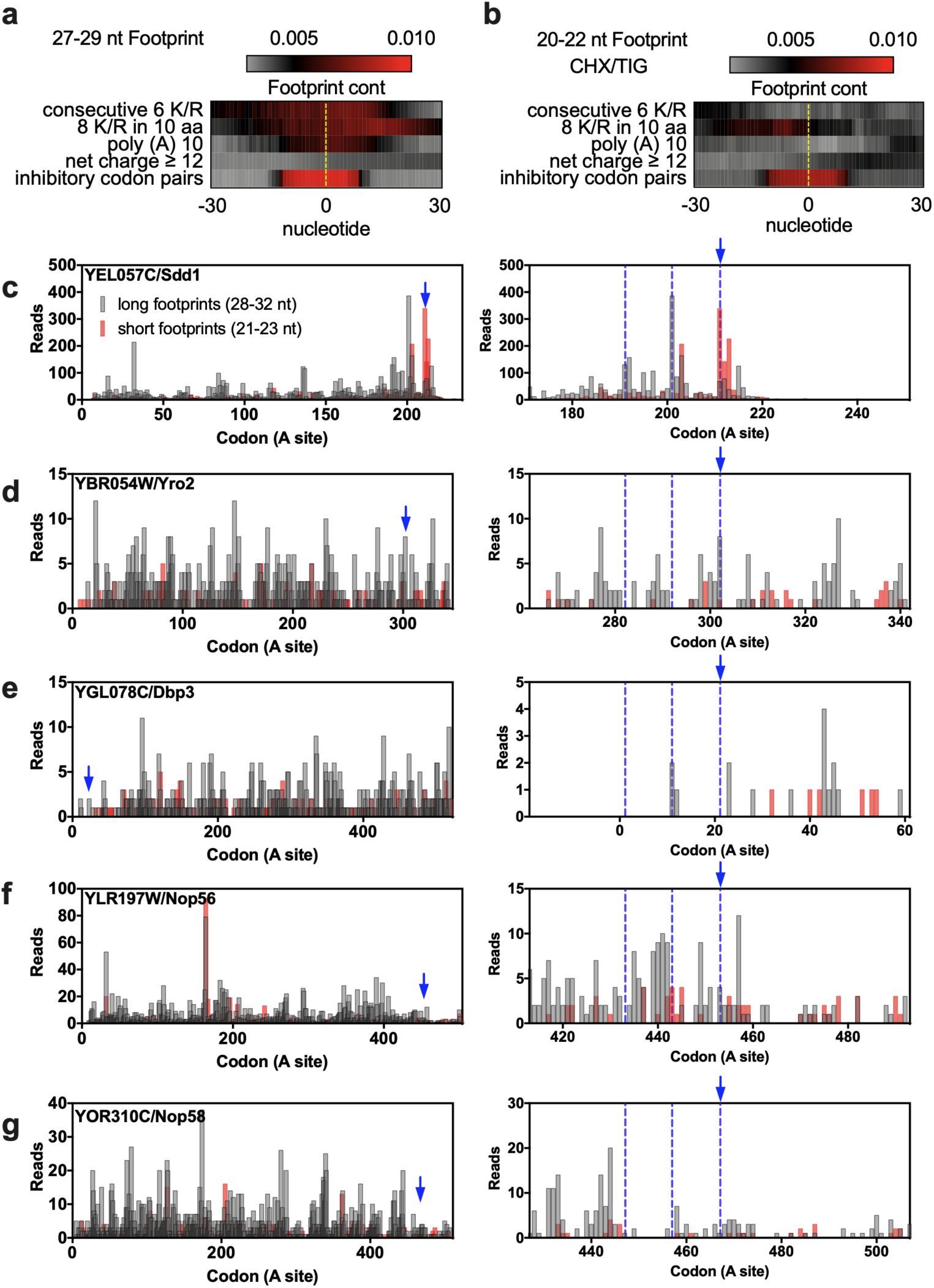
Ribosome profiling data of polybasic sequences. (a) 27-29 and (b) 20-22 nucleotide footprint analyses of genes containing polybasic sequences. As control, genes with one of the 17 inhibitory codon pairs (ICP) were used. The dotted yellow line represents the start of the polybasic sequence or of the ICP. (c) 28-32 and 21-23 nucleotide footprint analysis of Sdd1 shows an accumulation of short reads at its polybasic site, indicating ribosome stalling, and accumulation of long reads upstream of the stalling site with a periodicity of roughly 10 codons, indicating ribosome collisions, which are highlighted on the right panel. Blue arrow indicates the start of the polybasic sequences, and the blue dotted line indicates the hypothetical position of the collided ribosomes. (d-g) 28-32 and 21-23 nucleotide footprint analysis of the proteins with the highest polybasic sequences (YHR131C and YNL143C were omitted because of their low RP coverage). For panels a-b, the full read was used while for panels c-g, just the ribosomal A site read was used. The reads were plotted at an approximate position of the ribosomal A site.

**Figure 4.**
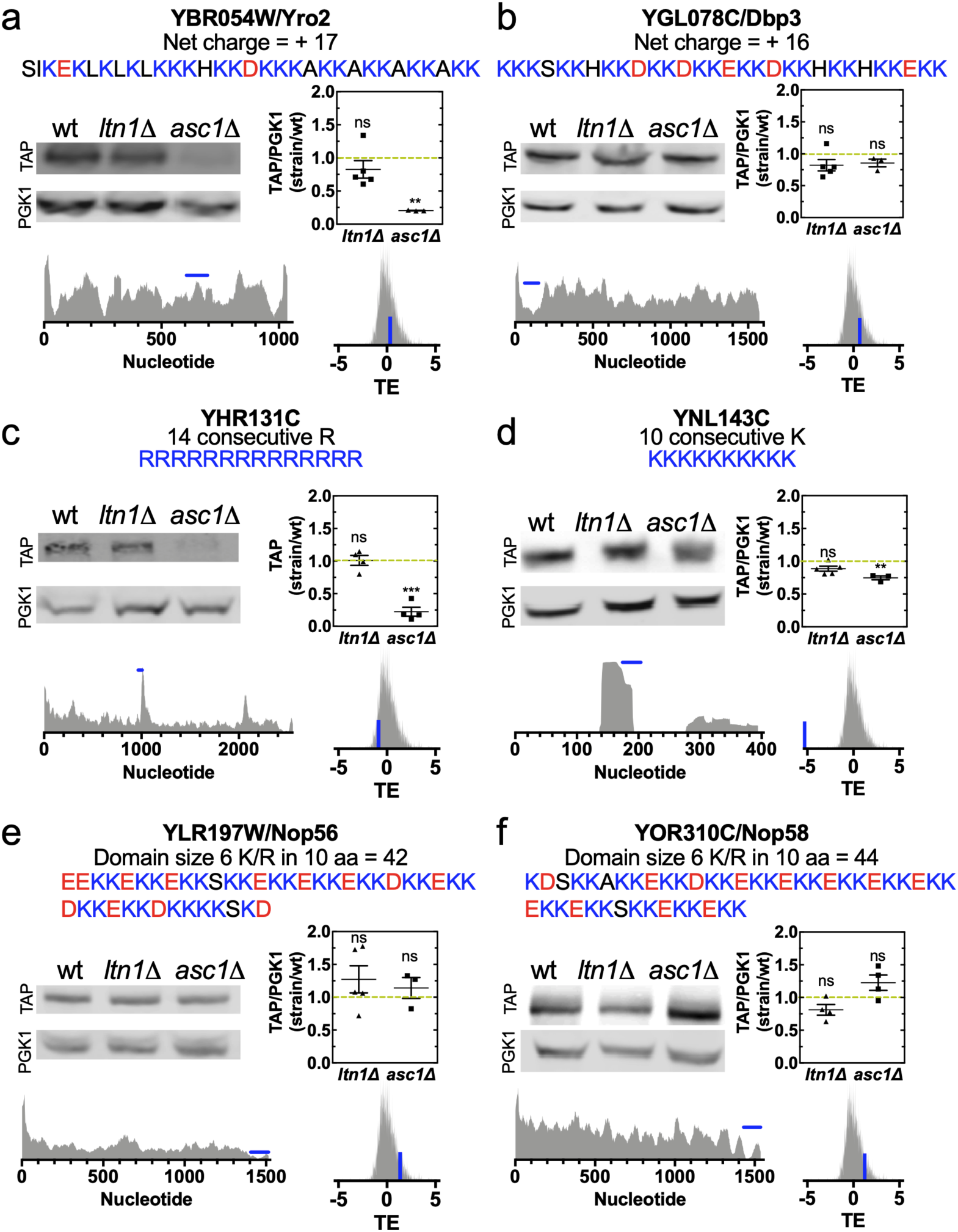
Proteins with the greatest polybasic sequences are not targets of the RQC complex in yeast. (a-f) Proteins with the greatest polybasic sequences of yeast were measured by western blot in wild type (wt), *lnt1*Δ, and *asc1*Δ strains. For each protein, the primary sequence of their polybasic sequence, their ribosome profiling data, and translation efficiency (TE) are presented. The blue bar in the ribosome profiling data indicates the position of the polybasic sequence, while the blue bar in the TE data shows the TE of the specific gene inside the TE distribution of all genes. The band intensity was quantified and normalized in relation to the housekeeping gene phosphoglycerate kinase (*PGK1*). One-way ANOVA with Bonferroni’s multiple comparison test was used for statistical analyses.

**Figure 5.**
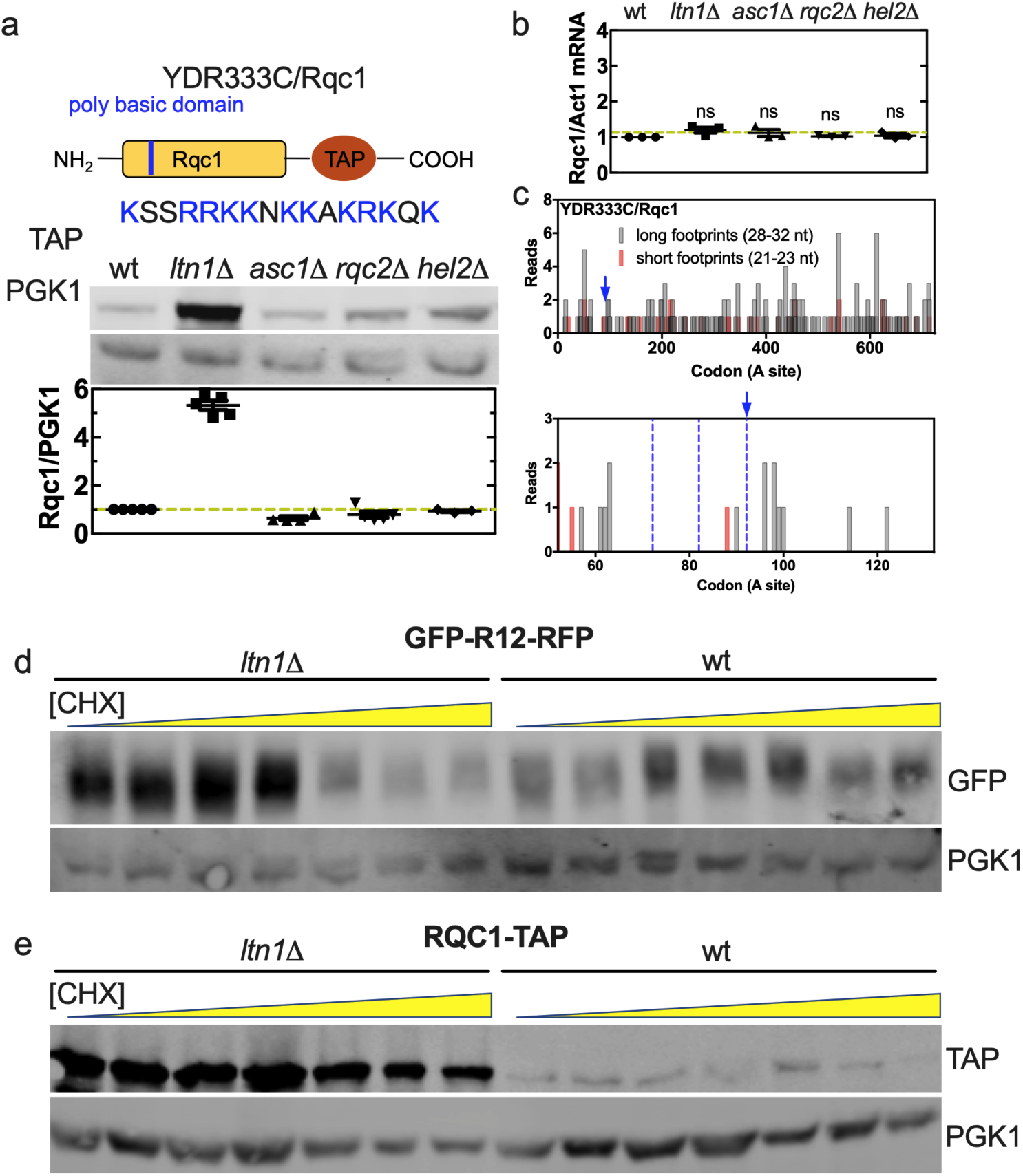
Rqc1 is not co-translationally degraded by the RQC complex. (a) A TAP-tagged version of Rqc1 was used to measure the protein levels in the wt, *lnt1*Δ, *asc1*Δ, *rqc2*Δ, and *hel2*Δ strains. The band intensity was quantified and normalized in relation to the housekeeping gene phosphoglycerate kinase (*PGK1*). (b) qPCR analysis of Rqc1-TAP mRNA levels in the wt, *lnt1*Δ, *asc1*Δ, *rqc2*Δ, and *hel2*Δ strains. The Rqc1 levels were normalized in relation to the housekeeping gene actin (*ACT1*). The analysis of differential expression was made by relative quantification using 2^−ΔΔ***Ct***^ method. One-way ANOVA with Bonferroni’s multiple comparison test was used for statistical analyses. (c) 28-32 and 21-23 nucleotide footprint analysis of Rqc1 shows neither accumulation of short reads at its polybasic site nor accumulation of long reads upstream of the stalling site (left and right panels, respectively). Blue arrow indicates the start of the polybasic sequences, and the blue dotted line indicates the hypothetical position of the collided ribosomes. The reads were plotted at an approximate position of the ribosomal A site. (d) Levels of the polybasic reporter GFP-R12-RFP subjected to 10 hours cycloheximide (CHX) treatment at increasing concentrations. (e) The same experiment as panel d was performed with a Rqc1-TAP strain.

### Read Length Distribution

For all datasets of ribosome profiling used herein, the read length distribution was calculated alongside the A-site prediction, using our own algorithm described in the Materials and Methods table.

### Clustering analysis

The clustering analysis of Figure 2c was performed by the Euclidean distance using Orange 3 software [51].

### Translation initiation normalized score

For Figure 2c, the genome was divided into eight equal groups based on their absolute values, from highest to lowest. The groups with the highest and the lowest values were further divided, making a total of ten groups. Each group was normalized individually. Their normalized values were then adjusted to fit the whole genome, meaning that the percentages from the group with the lowest values went from 0-100% to 0-6.25%, and the percentages from the group with the highest values, from 0-100% to 93.75-100%, for example. This methodology was used to avoid the influence of a few genes with extremely high and low values that can make the visualization of the difference between the values of most proteins difficult.

## RESULTS

### Identification of polybasic sequences in the proteome of *Saccharomyces cerevisiae*

Our first step was to find endogenous yeast proteins containing sequences with different arrangements of polybasic sequences. We selected four distinct, but possibly overlapping architectures, covering most of polybasic sequence variations analyzed in previous studies: (a) sequences of 8 or more lysines/arginines residues (K/R) in a window of 10 amino acids, (b) 6 or more consecutive K/R, (c) adenine repeats longer than 10 nucleotide residues, and (d) net charge ≥ +12 in a window of 30 residues. The architectures a) and b) were selected because they are often present in RQC activation reporters and lead to the accumulation of reads in ribosome profiling data [6, 9, 21-24, 26, 28, 49, 52-59]. The architecture c) was selected because poly (A) mRNA composition can lead to ribosome sliding, worsening the translation efficiency of these sequences [60, 61]. Architecture d) was selected because proteins containing a stretch of 30 residues (approximately the polypeptide length enclosed by the ribosome exit tunnel) with net charge ≥ +12 are extremely rare in most proteomes [40, 41]. A complete description of the selection of the architecture and cut-offs used are provided in Supplementary Material. We also compared these architectures with a group of 1800 genes containing inhibitory codon pairs (ICP) that are known to impair translation with substantial down-regulation in protein output (e.g., CGA-CGA and CGA-CCG) [62]. It is important to note that the ICP group, used herein as an endogenous positive control for ribosome stalling, has some potential pitfalls. For example, the sequence CGA-CCG-A, captured in our list of ICP as CGA-CCG (Supplementary Table S4), likely induces frameshifting [63]. Even though the pair CGA-CCG is present in just 1% of the genes in our list, we cannot rule out that other ICP arrangements might cause frameshifting and lead to poor translation scores by mechanisms unrelated to ribosome stalling.

We found that 354 proteins (approximately 6% of the yeast proteome) contained at least one of the characteristics described above (Figure 1a). The features most commonly found were high net charges and poly (A) sequences, with 170 and 121 sequences, respectively. Moreover, the Venn diagram (Figure 1a) shows that just 91 proteins have more than one feature at the same time, and only 1 protein, the uncharacterized YNL143C, has all the characteristics. Localization analysis revealed that no bias was found for the different categories analyzed (Figure 1b). We selected YNL143C and other proteins with prominent polybasic sequences (Figures 1c-e) for further experimental analysis in the next sections.

### Proteins with polybasic sequences show divergent patterns of translation efficiency parameters

If the presence of polybasic sequences represents an important obstacle to translation, we expect to find signatures of poor translatability in proteins that contain them. For example, endogenous genes containing stall sequences have a slower or less efficient translation initiation; this is thought to be an adaptation to avoid ribosome collision during elongation [32]. Therefore, we asked whether the polybasic and the ICP data sets would be significantly different from the rest of the yeast genes in experimental measurements that reflect translation activity. We chose six parameters that indicate how well a transcript is translated: (1) translation efficiency (TE) derived from ribosome profiling data, calculated from the ratio of ribosome-protected footprint read counts to the total mRNA read counts for each gene [64]; (2) codon stabilization coefficient (CSC), derived from mRNA half-life that positively correlates with TE and protein abundance [43]; (3) time for translation initiation, derived from ribosome profiling density, relative mRNA concentration and time of translation for individual codons [65]; (4) Kozak score 50 nt, determined by reporter expression quantification from a library containing 5’ 50 nt UTRs sequences from yeast genes cloned upstream the reporter’s start codon [66]; (5) Kozak score, determined by reporter expression quantification from a library containing synthetic 5 nt sequences cloned upstream the reporter’s start codon [66]; (6) 5’ UTR footprint density of the 40S ribosomal small subunits [67]. The Supplementary Table S3 shows how these metrics correlate with each other for the entire genome data-set. TE had a positive correlation with CSC, and both Kozak scores and negative correlation with time for translation initiation and 40S 5’ UTR densities. Moreover, even though we are using different ribosome profiling experiments to obtain the TE and time for initiation values, we still observed the expected positive correlation between these parameters (Supplementary Table S3).

In Figure 2a, we compare the distribution obtained with the entire genome with the distribution of values obtained with the ICP and polybasic gene groups. Consistent with the pattern expected for hard-to-translate sequences, the ICP group showed higher initiation times and 40S 5’ UTR densities, and lower TE, CSC, and Kozak scores. All these differences were supported by a t-test analysis comparing the individual data sets with the rest of the yeast genes (p-values are shown in Figure 2b). For the polybasic group, the only differences with a significant p-value were lower TE and CSC values than the genome group. Separating the polybasic sequences into sub-groups did not reveal a particularly deleterious architecture of polybasic sequence arrangement (Figure 2b). The cumulative distributions for each of the measurements described above are shown in Supplementary Figure S1.

To evaluate the characteristics of each gene from the polybasic group, we created a unified rank for all the measurements above. Using the values obtained with the entire yeast data set, each score system was organized in 10 bins; next, the maximum and minimum bin values were used to normalize the scale. We created a heat map (Figure 2c, upper panel) where 1 refers to the maximum translation activity (high TE, high CSC) or optimal initiation (short translation initiation time, high Kozak’s efficiency, and low 5’UTR footprint density of the 40S ribosomal small subunits). A cluster analysis of the 354 genes with polybasic sequences revealed that the group is rather heterogeneous and displays all spectra of characteristics, with some sequences having excellent translatability indicators with poor initiation scores and vice versa (Figure 2c). The lower panel indicates the polybasic architecture present in each gene. Notably, the sequences with the highest positively charged sequences or longest poly (A) tracts also showed diverse features (Figure 2c, lower panel). Therefore, we could not find a clear signature of poor translatability for genes containing polybasic sequences using the analyzed parameters (Figure 2c-d).

### Ribosome profiling of endogenous proteins with polybasic sequences

To test if the selected polybasic sequences have any effect on translation, we took advantage of previously published ribosome footprint profiling data. Analyses of the ribosome profiling (RP) data of the polybasic and ICP sequences indicated an enrichment of 27-29 nucleotide long footprints around the staling sequence (Figure 3a). We also plotted the number of reads at an approximate position of the ribosomal A site and observed an accumulation of reads downstream the first codon of the polybasic sequence (Supplementary Figure S2), which suggests a reduction in translational speed.

Additionally, we looked directly at the presence of ribosome collisions through the analyses of disome profiling. The disome profiling allows sequencing of mRNA fragments protected by two stacked ribosomes and shows stalling patterns undetected by traditional RP [29, 36, 37]. Disome footprints could be detected as a sharp peak immediately after the ICPs sequences (Supplementary Figure S3a-c). The analysis of polybasic sequences also showed an increase in the disome footprints. However, the peak was broader than the one observed with ICPs and occurred 9-10 codons after the polybasic-coding nucleotides (Supplementary Figure S3d-f). This observation supports the hypothesis that the arrest is caused by electrostatic interactions of the nascent positively charged polypeptide with the ribosome exit tunnel. Since disomes were detected in most transcripts, it is unlikely that the formation of these disomes can trigger the RQC activation [29, 37].

Recently, it was shown that ribosomes with open A sites produce shorter mRNA fragments (20-22 nucleotides) than ribosomes with occupied A sites, which produce the longer mRNA fragments (27-29 nucleotides) analyzed in Figure 3a [35]. Structural and biochemical evidence demonstrated that the electrostatic repulsion caused by the presence of a polybasic sequence inside the exit tunnel can decrease the binding efficiency of an incoming tRNA carrying an additional positively charged residue, leading to empty A site ribosome [25, 39, 68]. Since ribosomes with empty A-site are recognized by translation termination factors, the analysis of 20-22 fragments could more accurately reflect translation efficiency [39]. The ICP was the only group that showed an accumulation of short reads associated to empty A sites (Figure 3b). The same result was observed with another RP dataset obtained with a different combination of translation inhibitors (cycloheximide and anisomycin) (Supplementary Figure S4). These results suggested that polybasic sequences can reduce elongation rates but they are not enough to cause severe ribosome stalling leading to the accumulation of open-A site ribosomes.

Another indicative of severe ribosome stalling is the occurrence of ribosome collisions upstream an empty A-site ribosome [26, 28]. If the endogenous polybasic sequences are strong stalling sequences, we should observe not only an increase in short nucleotide footprints corresponding to an empty A-site, but also an increase in long nucleotide footprints upstream of the stalling site with a periodicity of roughly 10 codons, which corresponds to the collided ribosomes [25]. Among the 354 genes with one or more polybasic sequences identified in our bioinformatics screening, we selected the four top genes (Figure 1c-e) for subsequent analyses: YBR054W/Yro2 (net charge = +17), YGL078C/Dbp3 (net charge = +16), YHR131C (14 consecutive arginines) and YNL143C (32 consecutive adenines coding for 10 lysines) (Figures 2 and 3). To expand our analyses, we also included genes with the two longest polybasic sequences, stretches with 6 or more K/R in 10 amino acids of the yeast proteome: YLR197W/Nop56 and YOR310C/Nop58 (42 and 44 residues long polybasic sequences, respectively). As a positive control, we included the gene *SDD*1, which has been recently described as an endogenous target of the RQC complex and shows the ribosome collision footprint pattern described above [25]. Our analyses revealed that, apart from *SDD*1, none of the selected genes had the periodic footprint pattern that would indicate ribosome collisions (indicated by dashed lines) (Figure 3c-g). A downside of this analysis was the low ribosome profiling coverage of the polybasic genes, and this is the reason why YHR131C and YNL143C were not included in Figure 3c-g. In an attempt to enhance the analysis sensitivity, we carefully examined all the biological replicates looking for consistent read patterns that could suggest the presence of queued ribosomes. We also checked the ribosome profiling of a *hel2*Δ strain, where the stalled ribosomes are more stable and produce stronger signals corresponding to the periodic pattern of collided ribosomes than in the wt strain (Supplementary Figures S5-9). *SDD*1 presented a reproducible pattern across all the replicates and a stronger queued ribosomes signal in the *hel2*Δ strain (Supplementary Figure S5). However, no consistent periodicity or queued ribosomes signal increase in the *hel2*Δ strain could be observed for the polybasic genes (Supplementary Figures S6-9). The ribosome collision footprint pattern was also carefully investigated for the remaining 348 proteins with polybasic sequences but, due to the low ribosome profiling coverage, no conclusion was reached (data not shown).

In conclusion, we observed that the ICP group of genes produced more features associated to ribosome stalling when than the polybasic group (Figure 3 and S3), therefore polybasic sequences may represent a superable obstacle to translation, and their presence in not enough to induce RQC activation.

### The six yeast proteins with most prominent polybasic sequences are not targets of the RQC complex

To confirm that polybasic sequences are not sufficient to cause ribosome stalling and translation termination, we examined whether the deletion of RQC components had any effect on the final protein output of our selected proteins, which would imply a regulatory role for the RQC complex on their expression. TAP-tagged versions of the 6 genes selected from the previous section were obtained from a yeast library containing a chromosomally integrated TAP-tag at the 3’ region of each gene, thus preserving the natural promoter region and the 5’ UTR of the transcripts [44]. We chose to delete Ltn1, the E3 ubiquitin ligase of the RQC, and Asc1, involved in the modulation of RQC activity because they represent different steps in the RQC pathway and have been consistently used in the investigation of its mechanism of action [6, 20, 25, 59, 69-71]. Each TAP-tag strain was then used to generate the *ltn1*Δ or *asc1*Δ variants, confirmed by PCR and western blot (Supplementary Figure S10). In Figure 4, we show the protein expression of each gene as probed by anti-TAP western blot in the 3 different backgrounds and the relative quantification of at least 3 independent experiments. For each gene, we also show their ribosome profiling (complete sequence, with the polybasic sequence marked in blue) and their TE (depicted as a blue bar relative to total distribution of TE values of the yeast genes). We observed that deletion of *LTN1* caused no change in the protein levels of the selected polybasic proteins. Since the TAP-tag is downstream the putative stalling sequence, the lack of Ltn1 should not stabilize the full-length construct. Although the results from the *ltn1*Δ strains may have a limited contribution for the understanding of the role of RQC in the regulation of the analyzed genes, it showed that these proteins are not regulated by Ltn1 in an RQC independent pathway, as observed for Rqc1 (see next section).

The results with *asc1*Δ revealed no changes in YGL078C/Dbp3, YNL197W/Nop56, and YOR310C/Nop58 protein levels, and a down-regulation in YBR054W/Yro2, YHR131C, and YNL143C (Figure 4). This pattern could not be explained by any feature, such as ribosome density on the polybasic site, TE, or initiation time. Asc1 has been linked to the assembly of the translation preinitiation complex, and its deletion impairs translation of a subset of genes, particularly small ORFs and genes linked to mitochondria function [72, 73]. This RQC unrelated roles of Asc1 could be responsible for the decreased levels since YBR054W/Yro2 protein is localized to the mitochondria and YNL143C protein is encoded by a small ORF.

Nevertheless, all tested proteins differed from what was observed with reporters containing stalling sequences, which had their expression levels up-regulated in *asc1*Δ strains [6, 20, 59, 69-71]. Therefore, we conclude that, even though Asc1 positively regulates the expression of some of the proteins, the complete RQC pathway is not actively degrading any of the endogenous proteins tested.

### Ltn1 regulates Rqc1, but the mechanism is independent of the RQC complex

As described above, the protein Rqc1 and Sdd1 have been previously reported as targets of the RQC complex. The regulation of Sdd1 shows the canonical pattern of RQC activity. Deletion of *LTN1* increased the levels of the truncated N-terminal of Sdd1, while deletions of *ASC*1 or *HEL2* increased the levels of its full-length protein [25]. On the other hand, the experimental data available on Rqc1 regulation showed increased levels of the full-length construct in a *ltn1*Δ background, when the expected would be the stabilization of the N-terminal fragment up to the stalling polybasic region [6]. This result is not consistent with a co-translational model of regulation Rqc1 regulation by RQC.

To investigate the mechanism of Rqc1 regulation by the RQC, we analyzed Rqc1 expression in the backgrounds *ltn1*Δ, *asc1*Δ, *rqc2*Δ, and *hel2*Δ. For this, we used a TAP-tagged version of Rqc1, meaning that we can only observe an increase in the levels of the full-length peptide. As previously described, the deletion of *LTN1* increased the expression levels of Rqc1, but, different from Brandman *et al.*, 2012, the deletion of *ASC1, HEL2* or *RQC2* had no impact on the full-length Rqc1 expression levels (Figure 5a). Rqc1 regulation was different from that of GFP-R12-RFP, an artificial reporter with a polybasic sequence known to be a target of the RQC [6, 20, 21, 23], which presented increased levels of GFP (truncated peptide) when *LTN1* was deleted, and increased levels of both GFP and RFP (full-length peptide) when *ASC1* or *HEL2* was deleted (Supplementary Figure S11). It is noteworthy that the Rqc1 polybasic sequence (blue arrow) is located at the protein N-terminal, meaning that only the upstream polypeptide fragment should be stabilized in a *ltn1*Δ strain. Therefore, the overexpression of the full-length Rqc1 in the *ltn1*Δ strain (Figure 5a and Brandman et al., 2012) is inconsistent with the current model for RQC action. The *RQC1* mRNA levels were the same in all tested backgrounds (Figure 5b). To test if the presence of the C-terminal TAP-TAG influenced *RQC1* mRNA levels, we repeated the qPCR analysis with the non-tagged WT *RQC1*, and the same results were observed (Supplementary Figure S12). Moreover, ribosome profiling of Rqc1 did not show the characteristic footprint pattern of ribosome collisions and accumulation of short fragments (Figure 5c – blue dashed lines, see also Supplementary Figure S13), further indicating that the Rqc1 polybasic sequence does not act as a strong stalling sequence. Another way to verify whether a transcript is a target of the RQC complex is through the saturation of the rescue system; if the protein of interest is a target, the saturation of the RQC will increase its expression levels. The saturation can be accomplished by treatment with low concentrations of an elongation inhibitory drug, such as cycloheximide (CHX), which leads to multiple events of ribosome stalling and RQC activation [6, 26]. Accordingly, Figure 5d shows that the GFP signal from the GFP-R12-RFP reporter increased at low CHX concentrations and that this effect was Ltn1 dependent (Figure 5d). However, when we tested the Rqc1-TAP-TAG strain, we could not observe the same result, further indicating that the RQC complex does not control the final Rqc1 protein output during translation (Figure 5e).

## DISCUSSION

The hypothesis that the translation of positive amino acids leads to ribosome stalling and translation repression has been disputed since shortly after its proposal. The Grayhack group was the first to propose that codon bias was the major factor behind ribosome stalling observed with positive sequences [70, 73-75]. They examined the influence of all sequences of ten repeated codons on luciferase activity, and they demonstrated that the codon choice had a stronger effect than the final charge. Of particular interest is the codon CGA, which leads to a 99% reduction in luciferase activity in comparison with the codon AGA, both coding for arginine [76]. In recent works, they proposed that the CGA codon does not need to be highly concentrated to cause translational repression: the presence of as few as two consecutive CGA codons [71, 75] or specific combinations of CGA with a second codon, the so-called inhibitory pairs [62, 77], are sufficient to inhibit translation. In the same token, the Green and Djuranovic groups proposed that poly (A) sequences are the major cause of translation repression observed with polylysine stretches [11, 39, 60, 78]. They showed that the presence of six AAA codons in tandem leads to an approximately 30% to 70% reduction in protein expression in comparison to AAG in different organisms and that this reduction is proportional to the quantity of AAA [11, 60]. Letzring *et al.*, 2010, had already observed that the translation of the codon AAA is worse than that of AAG [76], but what both the Green and the Djuranovic groups demonstrated is that the translation disruption is caused by the presence of a poly (A) sequence in the mRNA, independent of codon choice. Sequences of AAA codons followed or preceded by codons rich in adenine (preceded by CAA and followed by AAC, for example) can cause greater translation repression than the same sequences of AAA codons alone [11]. The presence of an extended poly (A) sequence can cause ribosomal sliding, a slow ribosome frameshifting movement that leads to the incorporation of extra lysine residues in the nascent peptide. More importantly, it leads to the translation of premature termination codons, which are targeted by nonsense-mediated decay and lead to translation termination [60]. This may be caused by the helical conformation that poly (A) sequences can adopt, which allows them to enter the ribosomal A site and inhibit translation [39].

On the other hand, there is still evidence that the charge itself can play a role in translation dynamics. Positively charged sequences, independent of codon choice, are translated at slower rates than negatively charged or non-charged sequences [41, 49], which is still considered a hallmark of ribosome stalling. Additionally, proteins with a high density of negatively charged amino acids in a 30-residue stretch are far more common than proteins with a high concentration of arginines and/or lysines in 30 residues, suggesting a negative selection pressure against long polybasic sequences [41, 79]. Moreover, even in the papers that highlight the effect of codon choice on the inhibitory effect observed with polybasic sequences, we can see evidence that charge alone is still a relevant factor for translation efficiency. Both Arthur *et al.*, 2015, and Koutmou *et al.*, 2015, show that the translation of twelve AGA codons leads to a reduction in protein amount of approximately 50% [11, 60]. Taken together, the literature suggests that the translation of rare codons (CGA) and poly (A) sequences are detrimental for translation but does not rule out the effect of positively charged segments as RQC complex activation factors.

Our search for endogenous targets of the RQC complex was motivated by the possibility of translation repression caused by positively charged segments, as described above. Rqc1 was the first example of an endogenous protein with co-translational regulation promoted by the RQC complex [6]. Our results show that polybasic sequences similar to Rqc1 are relatively common (Figure 1), and even though they are translated at slower rates (Figure 3a and S2) [49], it seems that this is not sufficient to cause a severe ribosome stalling with empty A site, as characterized by the stabilization of 20 to 22 nucleotide footprints [35, 39] (Figure 3b, S4). Consistent with our results that polybasic sequences do not cause premature termination, not even the most positively charged sequences could be identified as endogenous targets of the RQC complex (Figure 4). Additionally, our results corroborate recent findings showing that short polybasic sequences promote ribosome collisions and are enriched in the ribosome exit tunnel of the disomes [29]. The authors argue that the translation of these disomes is likely to resume without recruiting the RQC pathway and, they hypothesize that these polybasic-induced pauses (and collisions) can favor the co-translational folding of upstream sequences [29]. Recently, the yeast gene *SDD1* was identified as a natural target of RQC. The staling sequence contains a rare combination of inhibitory features: a stretch of bulky amino acids, followed by a polybasic sequence with a CGA-CGA ICP and a short poly (A) tract. The result is a combination of amino acid−specific inactivation of the peptidyl-transferase center at the 60S subunit and mRNA-specific obstruction of the decoding center at the 40S subunit [25]. It is important to note that mutations on amino acids upstream of the polybasic sequence were able to increase the levels of Sdd1 protein expression [25], revealing that the polybasic sequence of Sdd1 was necessary, but not sufficient, to cause translation repression. Agreeing with our data, it appears that, in most cases, the translation of polybasic sequences does not lead to severe ribosome stalling, but it can become troublesome when combined with additional characteristics. Future studies will help us to identify and understand how these characteristics affect the mechanisms of co-translational regulation.

Similar to what was observed with other proteins with polybasic sequences, we showed that the RQC complex does not inhibit Rqc1 translation. In our hands, *rqc2*Δ, *hel2*Δ, and *asc1*Δ did not significantly affect Rqc1 levels, which speak against an RQC complex-dependent regulation (Figure 5a). However, Rqc1 overexpression in a *ltn1*Δ strain agrees with the result from Brandman *et al.*, 2012, which, at the time, was interpreted as evidence that the Rqc1-Flag nascent protein was being targeted by the RQC complex. However, the deletion of *LTN1* alone is usually associated with the stabilization of translation intermediates (up to the stalling point), as we observed in the GFP-R12-RFP reporter (Supplementary Figure S11), not full-length constructs as observed with Rqc1-TAP. Taking our data into account, we speculate that the K-rich basic sequence can function as the ubiquitylation site involved in Rqc1 regulation. Since it was previously reported that Ltn1 could ubiquitylate substrates independently of the RQC complex [80, 81], we speculate that Ltn1 might regulate Rqc1 in a process that is uncoupled from Rqc1 translation.

## Supporting information

Support Information

## DATA AVAILABILITY

The algorithms used in this work are available in the GitHub repository (https://github.com/mhoyerm).

## ACKNOWLEDGMENT

We thank Dr. Wendy Gilbert for providing us with the Asc1 antibody and Dr. Onn Brandman for providing us with a GFP-R12-RFP stalling reporter. We thank Dr. Nicholas Guydosh for providing us with the detailed ribosome profiling methodology process of disome dataset. We thank Dr. Nicholas Ingolia for providing us with the TE values of yeast genome. We also thank Santiago Alonso and Ramon Cid for their technical support.

## CONFLICT OF INTEREST

The authors declare that they have no conflicts of interest with the contents of this article.

**Supplemental data for this article can be accessed on the publisher’s website.**

