## Supplementary material for "Rqc1 and other yeast proteins containing highly positively charged sequences are not targets of the RQC complex": Support Information

**Support Information for:**

### These authors contributed equally.

#### MATERIALS AND METHODS

List of primers and PCR condition.

| Primer | Sequence 5'-3' | annealing temperature (T)/extension time (Time) |
|---|---|---|
| Ltn1-F-deletion | TAAGCCATCAAAAAAAGTTCAAGCAATAGTTGGTTCT TAATGCGTACGCTGCAGGTCGAC | T = 54°C/Time = 3 min |
| Ltn1-R-deletion | AAATGTCGTACATTTATATGAAATTTATATGCGATAG TCTAATCGATGAATTCGAGCTCG | |
| Ltn1F1 | GGAGAGGTCTCCGGTTCGATTCC | T = 60°C/Time = 1 min |
| KanB | CTGCAGCGAGGAGCCGTAAT | |
| Ltn1F1 | GGAGAGGTCTCCGGTTCGATTCC | T = 56°C/Time = 1 min |
| Ltn1R1 | CAGTTTAGCATATATCTGCGACCAACA | |
| Ltn1C | TAACAAATATCCAAGTTAATGGCGT | T = 54°C/Time = 1 min |
| Ltn1D | GAACAATTGAGAGAATGAAAAAGGA | |
| KanF1 | TATGGAACTGCCTCGGTGAG | T = 54°C/Time = 1 min |
| Ltn1D | GAACAATTGAGAGAATGAAAAAGGA | |
| Asc1-F-deletion | CCAAAAAATCCTTATAACACACTAAAGTAAATAAAGT GAAAAATGCGTACGCTGCAGGTC | T = 54°C/Time = 3 min |
| Asc1-R-deletion | CTAGAAGATACATAAAAGAACAAATGAACTTTATACA TATTCTTAATCGATGAATTCGAG | |
| Asc1F1 | CCTGGCCATCTGTAGCCTTA | T = 54°C/Time = 1 min |
| KanB | CTGCAGCGAGGAGCCGTAAT | |
| Asc1F1 | CCTGGCCATCTGTAGCCTTA | T = 54°C/Time = 1 min |

| | | |
|---|---|---|
| Asc1R1 | TGGCAACATCCCATAATCTCAAGG | |
| Asc1F2 | GGACGGTGAAATTATGTTGTGG | T = 54°C/Time = 1 min |
| Asc1R2 | CGCAGCAAACAGAAAGCATAG | |
| KanF1 | TATGGAACTGCCTCGGTGAG | T = 54°C/Time = 1 min |
| Asc1R2 | CGCAGCAAACAGAAAGCATAG | |
| Hel2-F-deletion | TCGAAAAAATAGTGGCTATACTTCTTTTGAAGAATTAGGATGCGTACGCTGCAGGTCGAC | T = 54°C/Time = 3 min |
| Hel2-R-deletion | ATGCTATTGTCAGTTACAGGTTAGAAATATATTTCCAACTAATCGATGAATTCGAGCTCG | |
| Hel2A | AGTGACCTCGTTATACATATCCCTG | T = 54°C/Time = 1 min |
| KanB | CTGCAGCGAGGAGCCGTAAT | |
| Hel2A | AGTGACCTCGTTATACATATCCCTG | T = 54°C/Time = 1 min |
| Hel2B | AAAGTCACAAACTTGTTCATCCTTC | |
| Hel2C | TTGAAAAGTCTCAACCTACCTCAAC | T = 54°C/Time = 1 min |
| Hel2D | ATTTTGTTGGAGTTGTCTTATGAGC | |
| Rqc2-F-deletion | TTATCCGGTCTAAGAAGTCAGGCAGGCAAGAGATTAATAGCGATGCGTACGCTGCAGGTCGAC | T = 54°C/Time = 3 min |
| Rqc2-R-deletion | AAAATTATAATTGCTGTCTATTTTCTTTTCATCTCATATGATTTATCGATGAATTCGAGCTCG | |
| Rqc2A | TCTGTGGTTACGATTAAAATTGGAT | T = 54°C/Time = 1 min |
| KanB | CTGCAGCGAGGAGCCGTAAT | |
| Rqc2A | TCTGTGGTTACGATTAAAATTGGAT | T = 54°C/Time = 1 min |
| Rqc2B | CAATCGACAACAACATTGAGTTTAG | |
| Rqc2C | ACAAGTTAAAAGTAACAATCGCTGG | T = 54°C/Time = 1 min |
| Rqc2D | ATTGATAATGGTGTTATCCTCGAAA | |
| KanF1 | TATGGAACTGCCTCGGTGAG | T = 54°C/Time = 1 min |
| Rqc2D | ATTGATAATGGTGTTATCCTCGAAA | |

**Rationale for cut-offs of polybasic sequences architecture in *S. cerevisiae*.**

Sequences of 8 or more lysines/arginines residues (K/R) in a window of 10 amino acids:
In previous analyzes, the *S. cerevisiae* genome was screened for sequences with 6 or more lysines and/or arginines within any group of 10 amino acids (1,2) and 1,187 proteins with polybasic sequences were found, which comprised 19% of the yeast proteins. We used a similar analysis herein but now with a more stringent cutoff, 8 or more K/R within 10 amino acids and found 84 proteins (1.4 % of yeast proteome, 8 K/R in 10 aa, Fig. 1a).

6 or more consecutive K/R:
Since most of reporters were made with K/R in tandem we screened for polybasic sequences with 6 or more K/R's in tandem and found 89 genes (1.5 % of yeast proteome, 6 K/R consecutives, Fig. 1a, 1c). From these analyzes we also identified the genes with highest consecutive K/R sequences, YHR131C (14 consecutives R) and YNL143C (10 consecutives K) (Fig 1c). The Inada group showed that the addition of a sequence of lysine residues to the C-terminus of a GFP-His3 construct reduced its expression levels and that this reduction was dependent on the number of lysine residues added (3). While the insertion of four lysine causes no effect into protein expression, the insertion of six lysine residues reduced protein expression levels to 45% (3).

Net charge ≥ +12 in a window of 30 residues:
The ribosome exit tunnel houses approximately 30 amino acids residues of the nascent polypeptide (4) and, because is formed primarily of nucleic acids, their electrostatic potential is negative (4). We developed a program that is able to screen the sequence of a given protein and calculate the net charge of every 30-amino acid segment (K and R = +1 and D and E = -1) (5). We found 170 genes (2.87 % of the yeast proteome) with at least one sequence with net charge ≥+12 (net charge 12, Fig 1a) being YBR054W/Yro2 (net charge = + 17) and YGL078C/Dbp3 (net charge = + 16) the genes with a 30 amino acid sequence with the highest net charge of yeast proteome (Fig 1d). The number of 30 amino acid

sequence with net charge = -12 is four-fold higher when compared with net charge = + 12 in yeast proteome (5).

Adenine repeats longer than 10 nucleotide residues

It was showed that the codon composition of poly basic residues has an important impact in translation (6). For example, an insertion of poly lysine-encoding sequences of $AAG_{12}$ in a report gene reduced the protein production by 50% while for the sequence $AAA_{12}$ the reduction was 80% (6). The insertion of the sequence $AAA_{10}$ into reporter leads to truncated peptide product (6). Consecutive AAA-lysines followed by ribosome sliding could potentially activate the RQC complex. Using a cutoff of ≥10 consecutive adenine (A) we found 121 genes (2 % of the yeast genome) (poly (A) 10, Fig. 1a, 1e). The highest concentration of poly adenine was 32, present in the gene YNL143C (Fig 1e).

Using the four different criteriums described before, we identified 354 genes (6% of the yeast proteome) and 91 genes have at least two features in common at the same time.

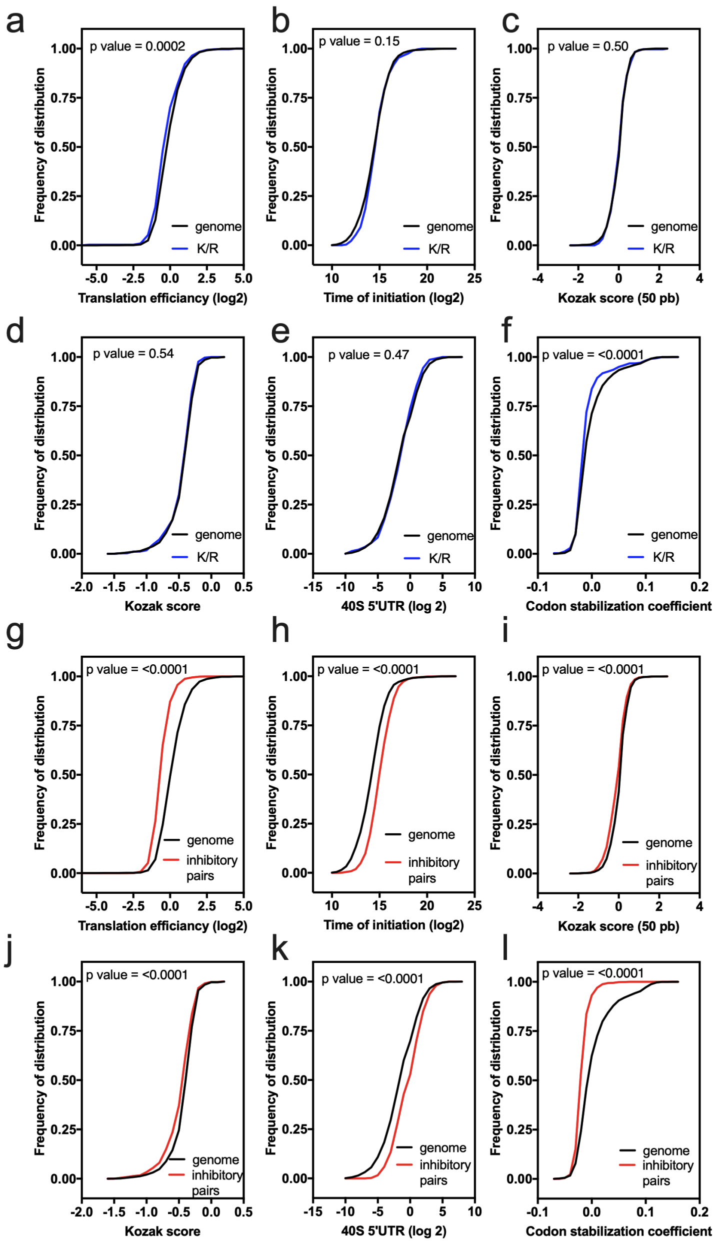


**Figure S1. Translation initiation rate of genes with polybasic sequences.** (a) The different groups of genes with some polybasic sequence were grouped in one single group (K/R) and compared regarding parameters that, to some extent, reflect translation initiation rate, with the genome (a-f). Kolmogorov-Smirnov test p values are showed for each comparison. As control was used, genes with one of the 17 inhibitory codon pairs (IP) (g-l). The parameters used to infer initiation gene rate were; translation efficiency, time for translation initiation, Kozak score of the 50 first nucleotide, Kozak score of the first 5 nucleotide, 40S ribosomal small subunit (SSU) footprint profiling (SSU 5'UTR: start codon) and codon stabilization coefficient.

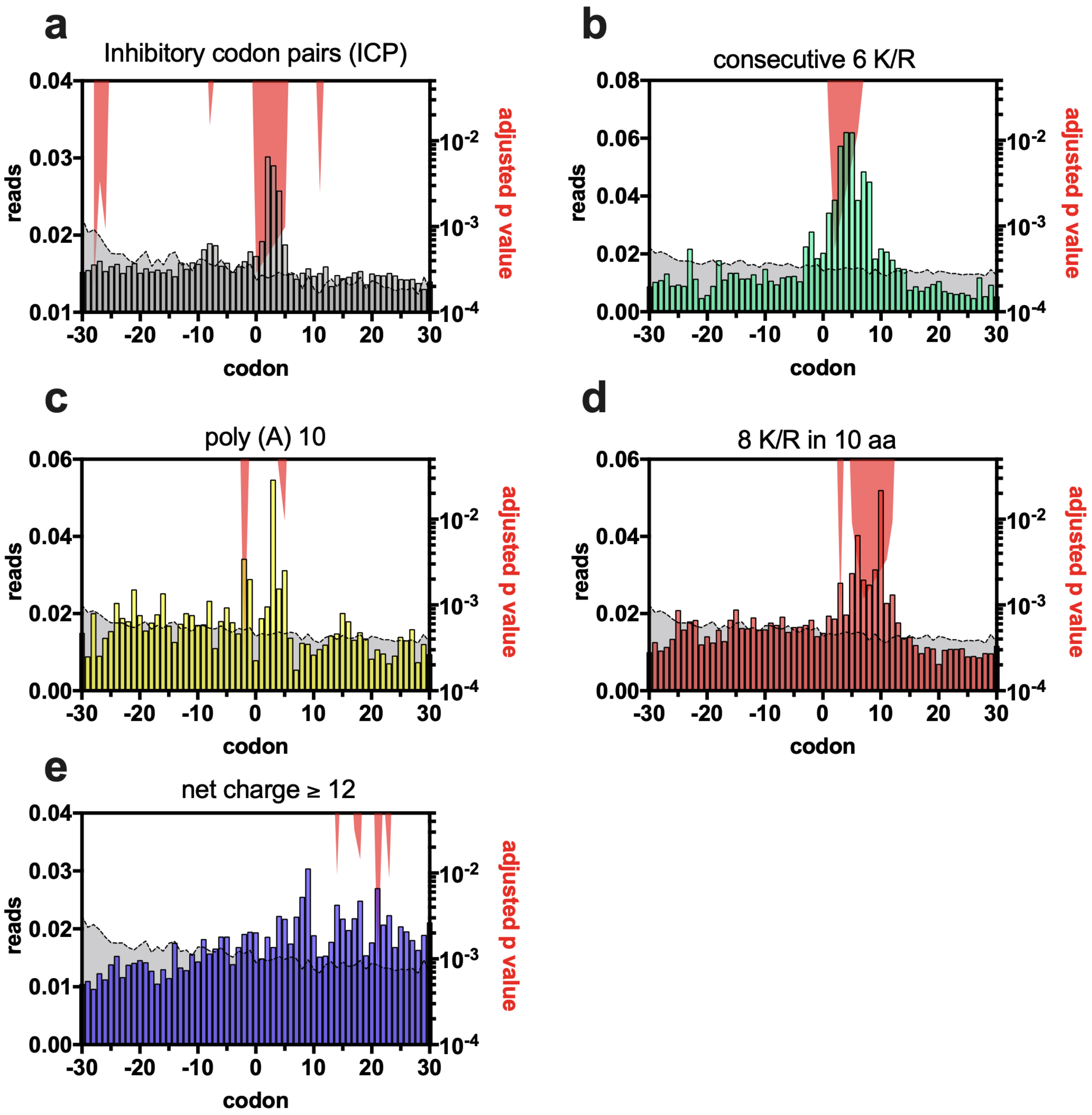


**Figure S2. A site ribosome profiling of genes with poly-basic sequences.** Ribosome profiling reads of the A site indicate that all poly-basic sequences are enriched with 28-32 nucleotide long footprints (b-e). As control, genes with one of the 17 inhibitory codon pairs (ICP) were used (a). The gray area represents the reads of a list of 2,000 random genes that allowed us to perform statistical analyses (multiple t-test using Holm-Sidak method, see details on methodology section). The right y-axis represents the adjusted p-value (red area).

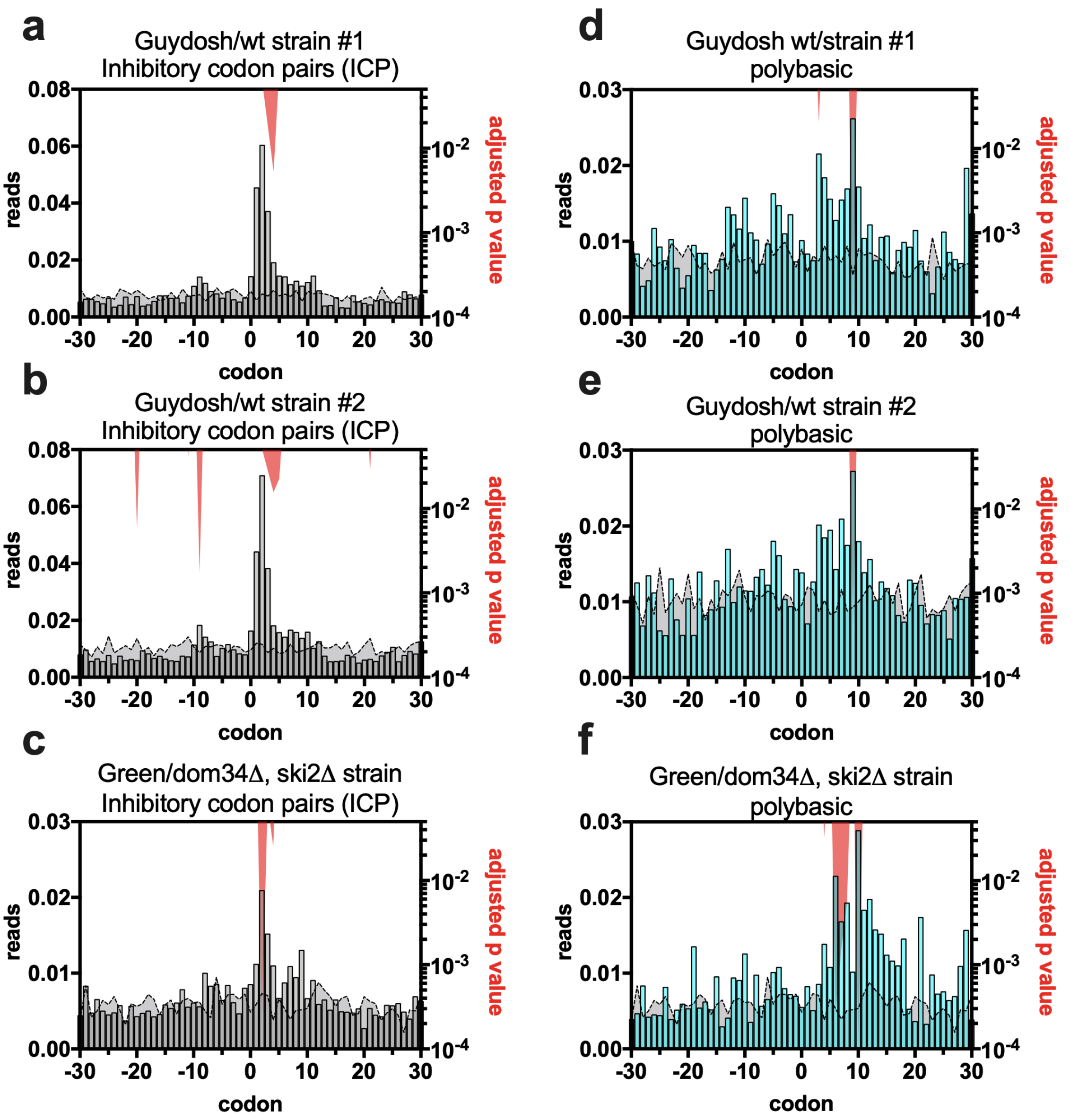


**Figure S3. Disome profiling of genes with poly-basic sequences.** Disome profiling reads of the A site indicate that genes with poly-basic sequences are enriched with disomes (d-f). As control, genes with one of the 17 inhibitory codon pairs (ICP) were used (a-c). The gray area represents the reads of a list of 2,000 random genes that allowed us to perform statistical analyses (multiple t-test using Holm-Sidak method, see details on methodology section). The right y-axis represents the adjusted p-value (red area). Three different disome datasets were used, two profiling's (#1 and #2) from a wild type strain (Guydosh: Meydam and Guydosh, 2020) and one profiling from a *dom34Δ, ski2Δ* strain (Green: D'Orazio et al, 2019). Due to the low ribosome profiling coverage, we grouped the four categories of polybasic sequences into one group namely, polybasic.

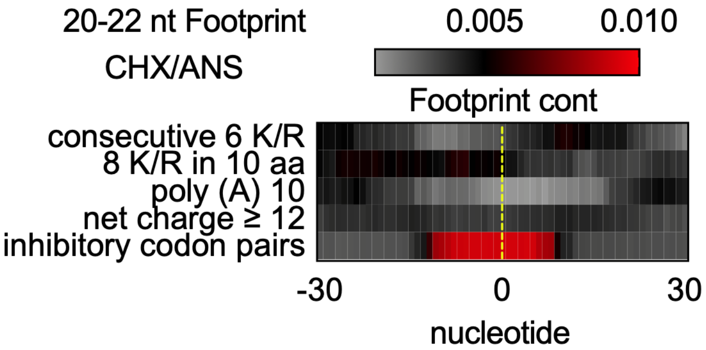


**Figure S4. Ribosome footprint profiling of the genes with polybasic sequences from yeast genome.** 20-22 nucleotide footprint analyzes of genes containing polybasic sequences. The dotted yellow line represents the start of the feature analyzed. As control was used, genes with one of the 17 inhibitory codon pairs (ICP). The 20-22 nt footprint was obtained by the use of cycloheximide and anisomycin (CHX/ANS).

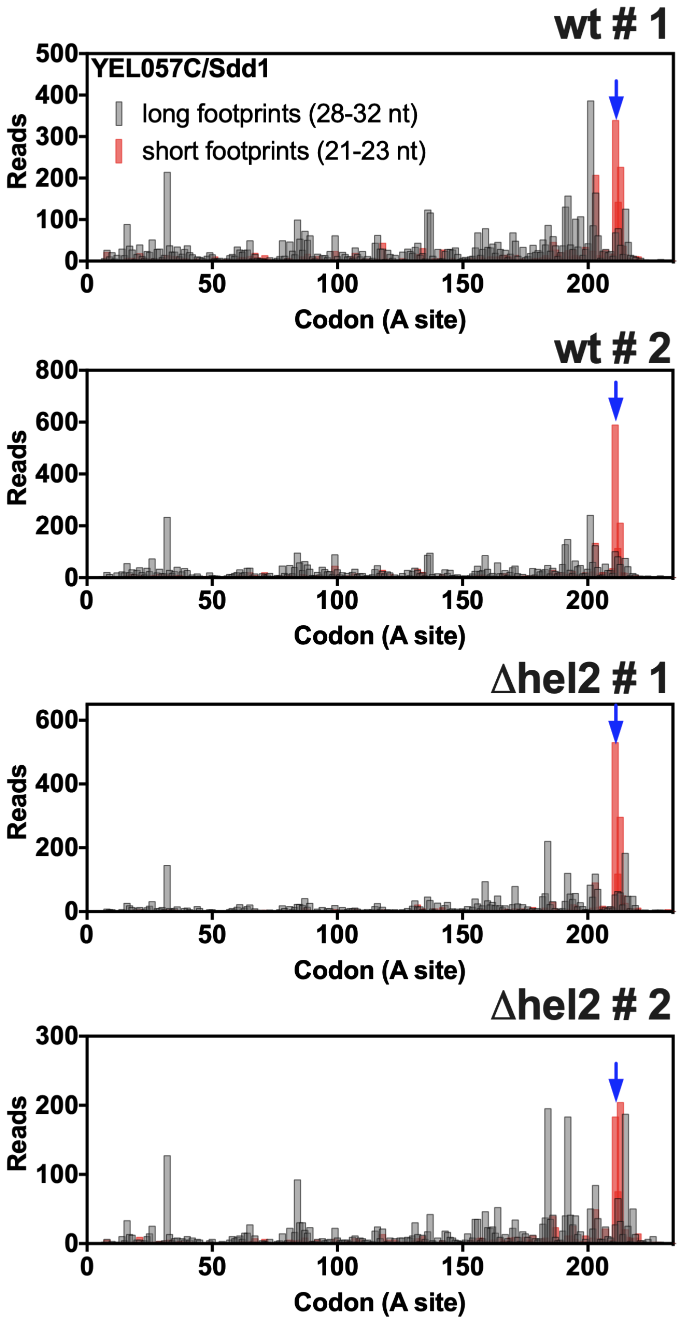


**Figure S5. Ribosome profiling data of *SDD1*.** 28-32 and 21-23 nucleotide footprint analysis of *SDD1* shows an accumulation of short reads at its polybasic site, indicating ribosome stalling, and accumulation of long reads upstream of the stalling site with a periodicity of roughly 10 codons, indicating ribosome collisions. The ribosome stalling is more evident in the *hel2Δ* strain. Blue arrow indicates the start of the polybasic sequences. The reads were plotted at an approximate position of the ribosomal A site. The two upper panels represent ribosome profiling obtained with a wt strain while the two in the bottom were obtained with a *hel2Δ* strain.

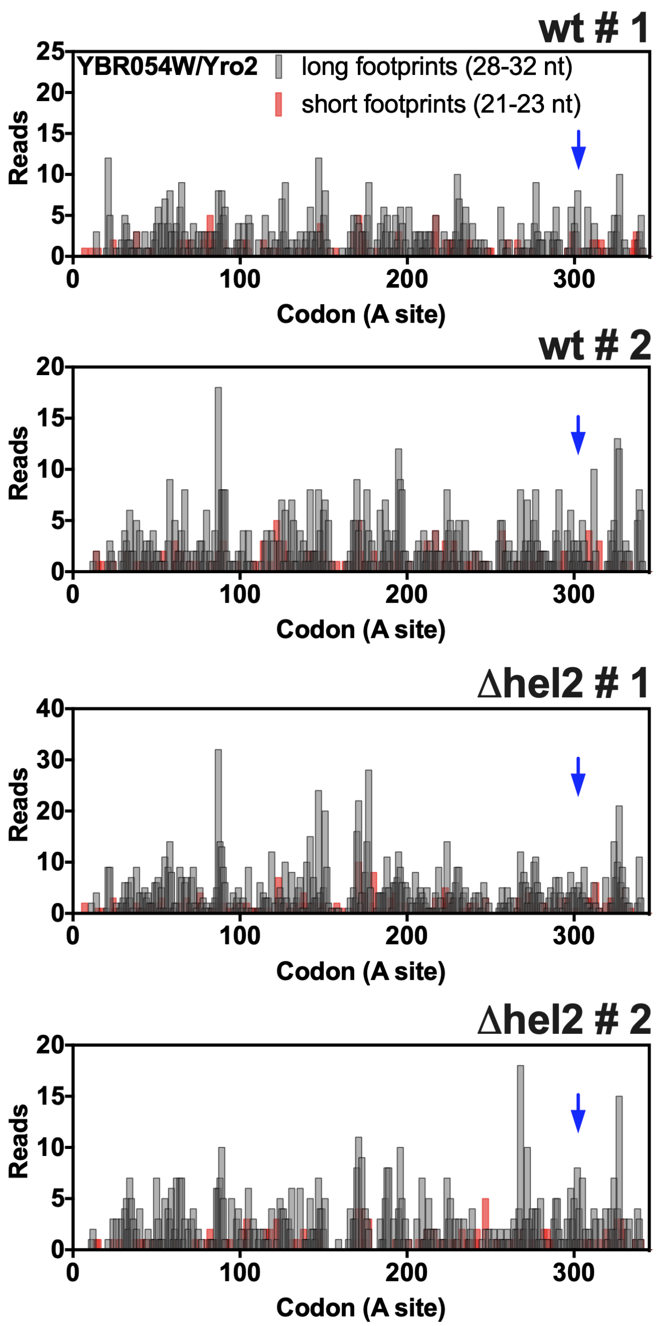


**Figure S6. Ribosome profiling data of *YRO2*.** 28-32 and 21-23 nucleotide footprint analysis of *YRO2*. Blue arrow indicates the start of the polybasic sequences. The reads were plotted at an approximate position of the ribosomal A site. The two upper panels represent ribosome profiling obtained with a wt strain while the two in the bottom were obtained with a *hel2Δ* strain.

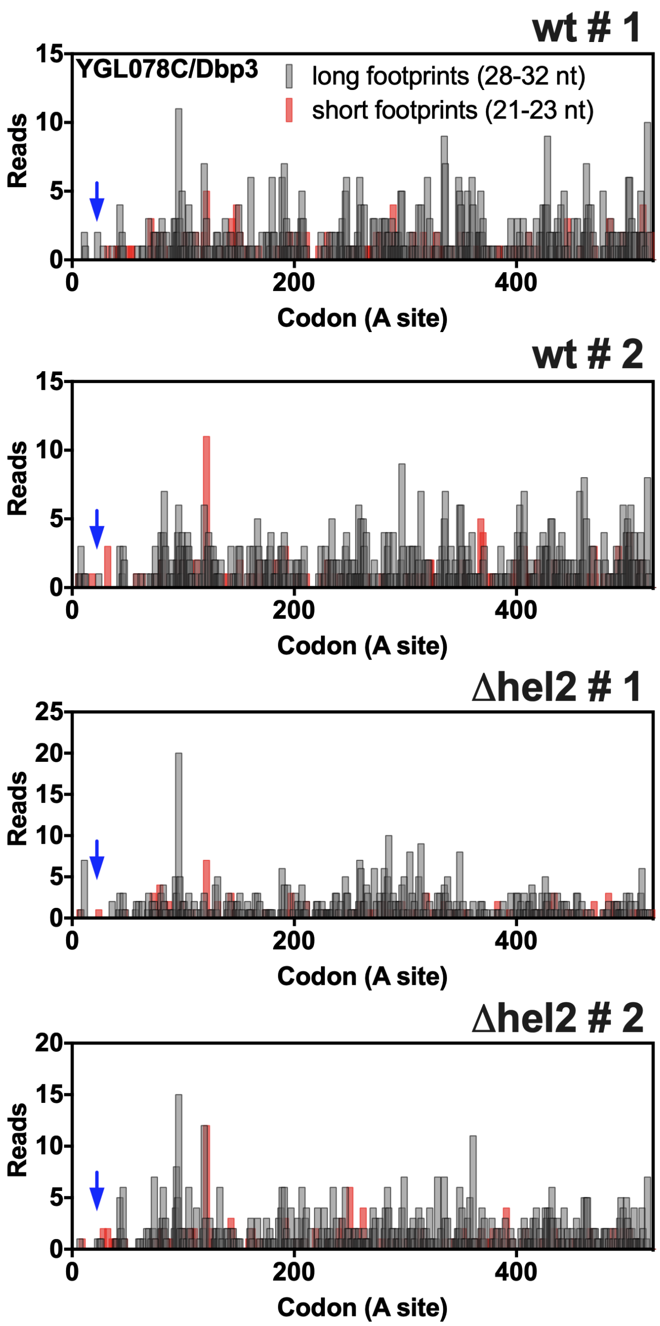


**Figure S7. Ribosome profiling data of *DBP3*.** 28-32 and 21-23 nucleotide footprint analysis of *DBP3*. Blue arrow indicates the start of the polybasic sequences. The reads were plotted at an approximate position of the ribosomal A site. The two upper panels represent ribosome profiling obtained with a wt strain while the two in the bottom were obtained with a *hel2Δ* strain.

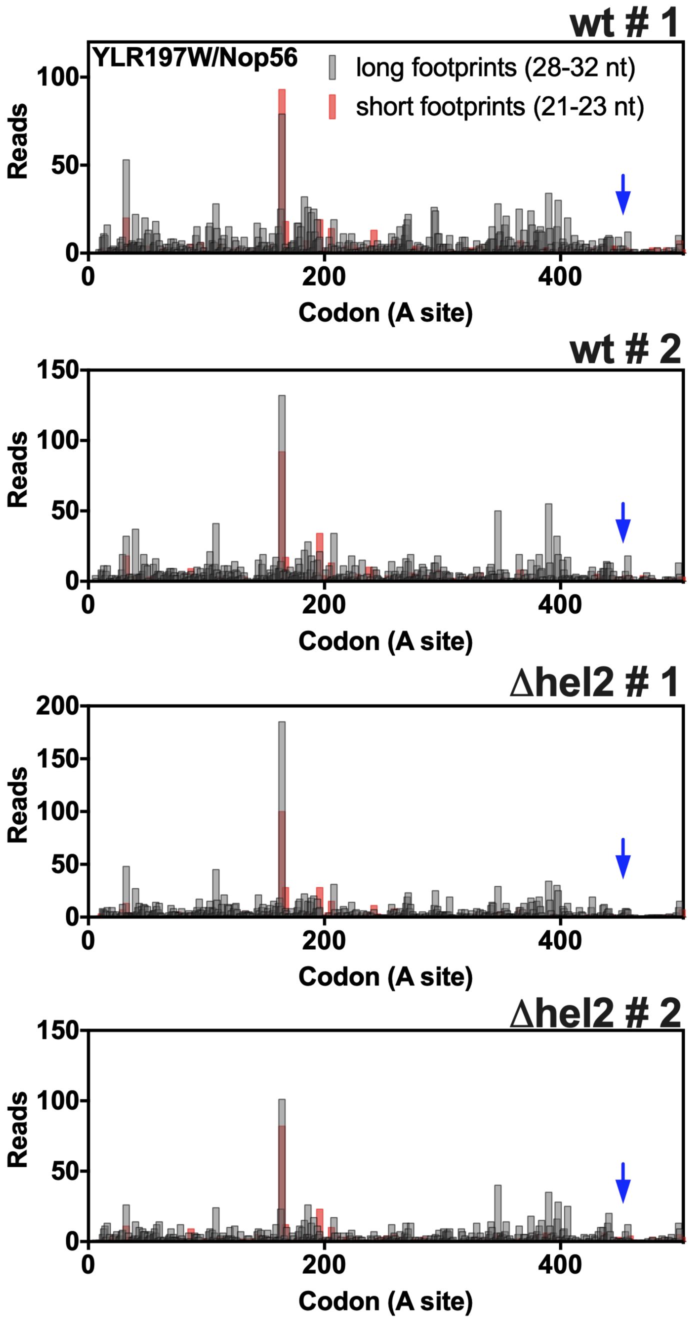


**Figure S8. Ribosome profiling data of *NOP56*.** 28-32 and 21-23 nucleotide footprint analysis of *NOP56*. Blue arrow indicates the start of the polybasic sequences. The reads were plotted at an approximate position of the ribosomal A site. The two upper panels represent ribosome profiling obtained with a wt strain while the two in the bottom were obtained with a *hel2Δ* strain.

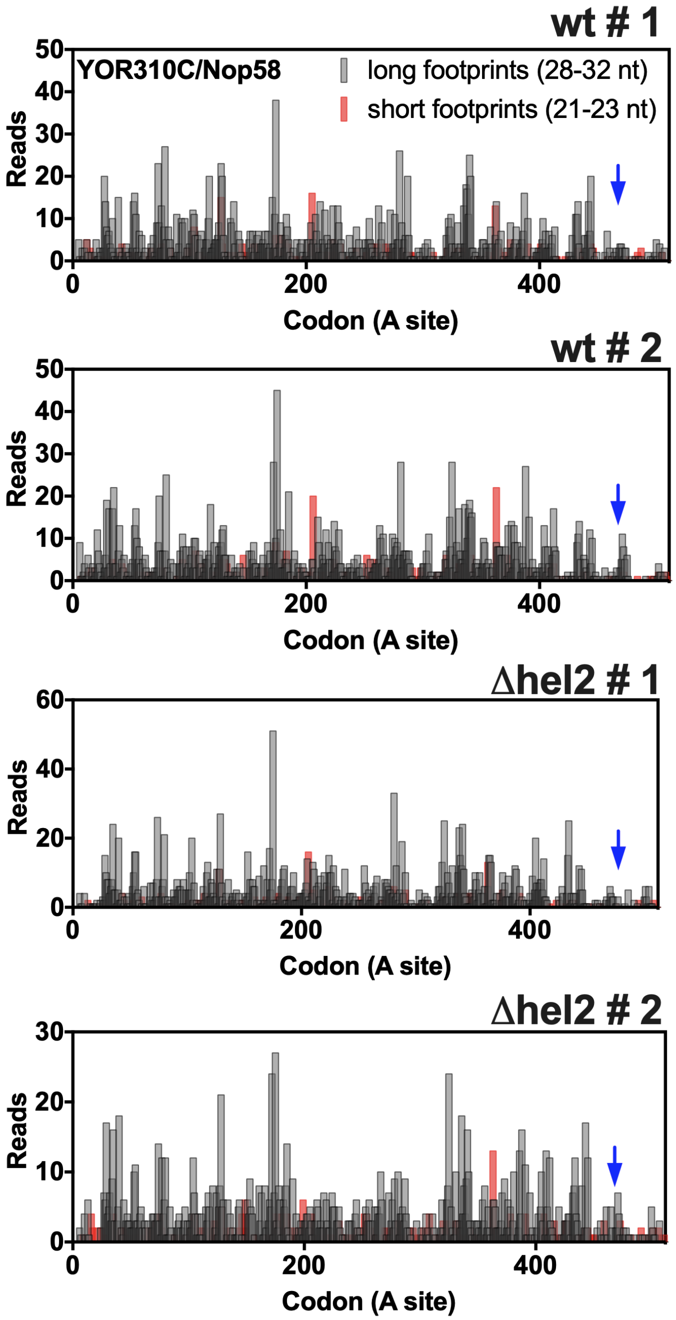


**Figure S9. Ribosome profiling data of *NOP58*.** 28-32 and 21-23 nucleotide footprint analysis of *NOP58*. Blue arrow indicates the start of the polybasic sequences. The reads were plotted at an approximate position of the ribosomal A site. The two upper panels represent ribosome profiling obtained with a wt strain while the two in the bottom were obtained with a *hel2Δ* strain.

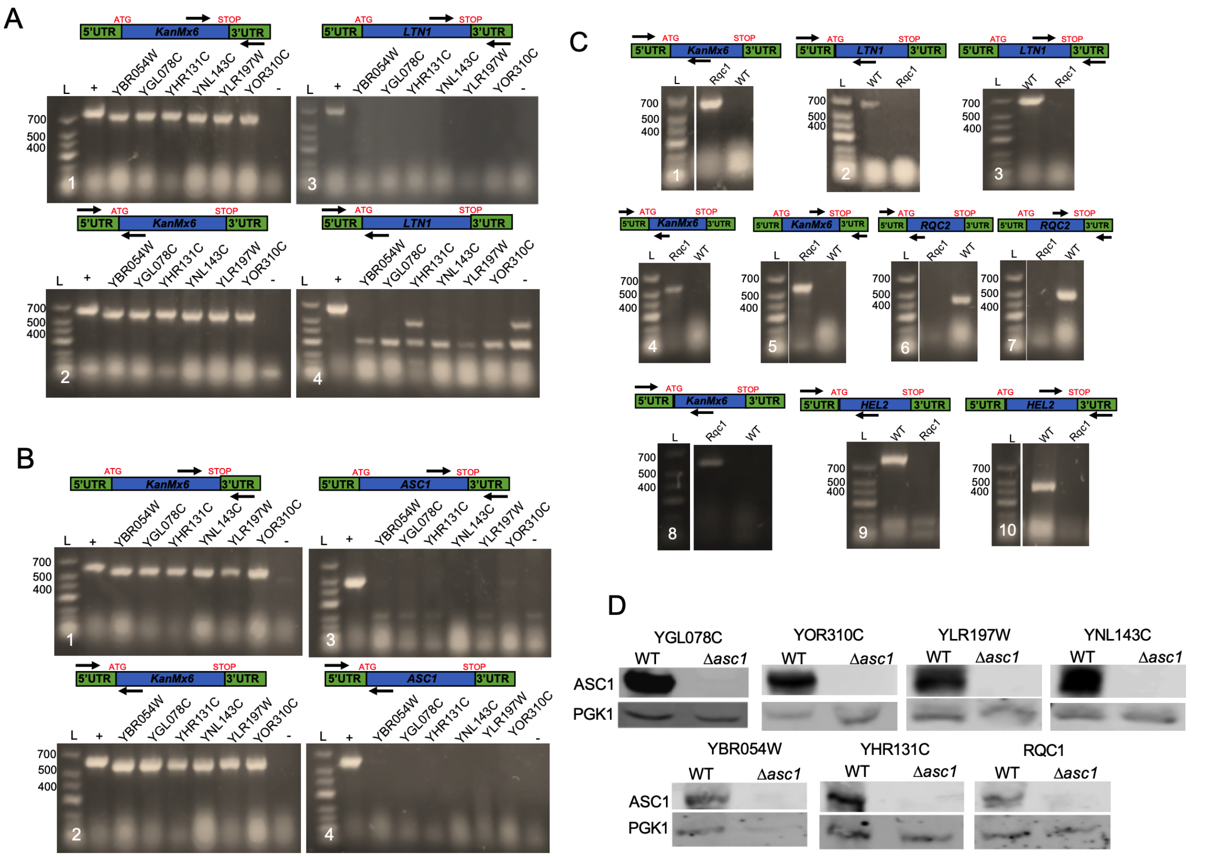


**Figure S10. Confirmation of deletions in the select TAP-TAG protein targets and Rqc1 TAP-TAG.** The gels showed in the panels a and b confirm the deletion of *LTN1* and *ASC1*, respectively, in all select targets. The four PCR sets show the presence of the *KanMx6* gene in the original gene ORF (gels 1 and 2), and the absence of *LTN1* (a) or *ASC1* (b) genes on their ORFs (gels 3 and 4). The DNAs from WT, *ltn1*Δ (knockout collection) and *asc1*Δ (knockout collection) strains were used as positive (+) or negative (-) controls of all the PCRs. (c) Deletions of *LTN1*, *RQC2* and *HEL2* were confirmed by PCR in the Rqc1-TAP TAG strain. For *LTN1* deletion, gel 1 shows the presence of *KanMx6* gene and gels 2 and 3 the absence of *LTN1* on its original ORF. For *RQC2* deletion, gels 4 and 5 show the presence of *KanMx6* gene and gels 6 and 7 the absence of *RQC2* on its original ORF. For *HEL2* deletion, gel 8 shows the presence of *KanMx6* gene and gels 9 and 10 the absence of *HEL2* on its original ORF. (d) Deletion of Asc1 in all targets and Rqc1 was confirmed by Western Blot using an anti-Asc1 antibody. L= ladder.

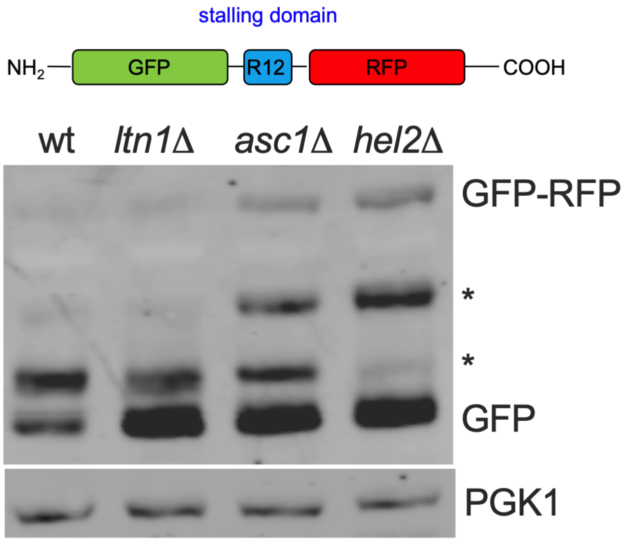


**Figure S11.** Whole-cell immunoblots of the indicated yeast strains expressing the stalling reporter (GFP-R12-RFP). GFP-RFP indicates the full-length translation product, and GFP indicates the arrest product. Stars denote products of the stalling reporter that likely to result from proteolytic cleavage.

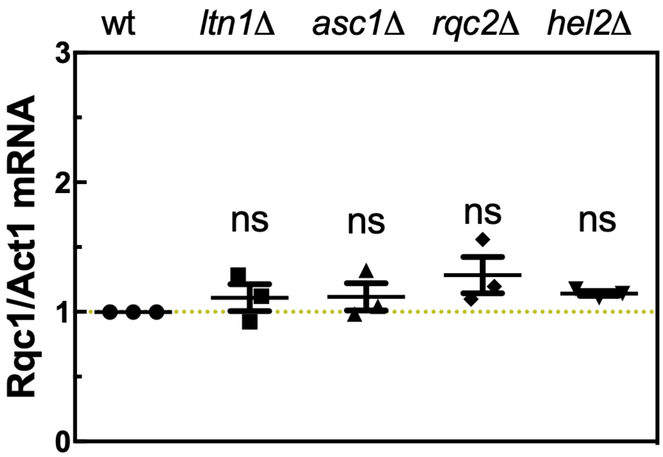


**Figure S12. Non-tagged Rqc1 mRNA levels**. qPCR was used to measure the non-tagged Rqc1 mRNA levels of the wt, *lnt1*Δ, *asc1*Δ, *rqc2*Δ, and *hel2*Δ strains. The Rqc1 levels were normalized in relation to the housekeeping gene actin (Act1). The analysis of differential expression was made by relative quantification using $2^{-\Delta\Delta Ct}$ method. One-way ANOVA with Bonferroni's multiple comparison test was used for statistical analyses.

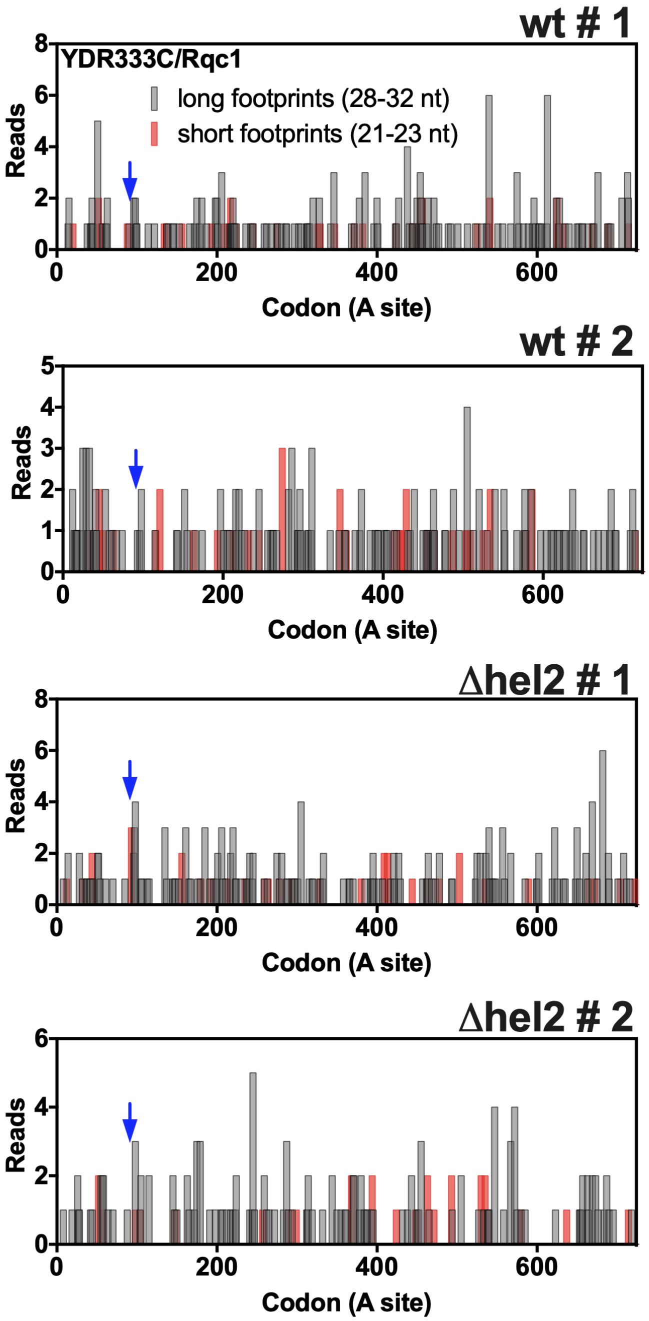


**Figure S13. Ribosome profiling data of *RQC1*.** 28-32 and 21-23 nucleotide footprint analysis of *RQC1*. Blue arrow indicates the start of the polybasic sequences. The reads were plotted at an approximate position of the ribosomal A site. The two upper panels represent ribosome profiling obtained with a wt strain while the two in the bottom were obtained with a *hel2Δ* strain.
